# Intracellular competition shapes plasmid population dynamics

**DOI:** 10.1101/2025.02.19.639193

**Authors:** Fernando Rossine, Carlos Sanchez, Daniel Eaton, Johan Paulsson, Michael Baym

## Abstract

Conflicts between levels of biological organization are central to evolution, from populations of multicellular organisms to selfish genetic elements in microbes. Plasmids are extrachromosomal, self-replicating genetic elements that underlie much of the evolutionary flexibility of bacteria. Evolving plasmids face selective pressures on their hosts, but also compete within the cell for replication, making them an ideal system for studying the joint dynamics of multilevel selection. While theory indicates that within-cell selection should matter for plasmid evolution, experimental measurement of within-cell plasmid fitness and its consequences has remained elusive. Here we measure the within-cell fitness of competing plasmids and characterize drift and selective dynamics. We achieve this by the controlled splitting of synthetic plasmid dimers to create balanced competition experiments. We find that incompatible plasmids co-occur for longer than expected due to methylation-based plasmid eclipsing. During this period of co-occurrence, less transcriptionally active plasmids display a within-cell selective advantage over their competing plasmids, leading to preferential fixation of silent plasmids. When the transcribed gene is beneficial to the cell, for example an antibiotic resistance gene, there is a cell-plasmid fitness tradeoff mediated by the dominance of the beneficial trait. Surprisingly, more dominant plasmid-encoded traits are less likely to fix but more likely to initially invade than less dominant traits. Taken together, our results show that plasmid evolution is driven by dynamics at two levels, with a transient, but critical, contribution of within-cell fitness.

## Introduction

Conflicts between levels of biological organization—genes, cells, organisms, populations—are an inescapable constraint of evolution[1]. These conflicts have shaped living beings from their emergence, when self-replicating molecules were first encapsulated in protocells[2], to the breakdown of cell-cycle control that leads to cancer and death in multicellular organisms[3]. Evidence of these conflicts has been found in fields as wide-ranging as experimental evolution[4–6], animal behavior[7], development[8], and molecular genetics[9]. Additionally, theory has predicted that these cross-scale processes can give rise to complex, non-intuitive evolutionary dynamics[10, 11]. Nonetheless, our empirical understanding of these dynamics lags behind the theory due to the general difficulty in simultaneously tracking the fitness of replicators across different levels of organization. In this context, plasmids—pervasive[12–15] extrachromosomal, self-replicating genetic elements that underlie much of the evolutionary flexibility of their bacterial hosts[16]—provide an ideal model system to study the dynamics of cross-scale conflicts.

Novel plasmid variants constantly emerge—oftentimes acquiring genes related to catabolism, virulence[17, 18], or antibiotic resistance[15, 19, 20] amongst other functions—and spread across bacterial communities[21], a process most notably exemplified by repeated selective sweeps of antibiotic resistance plasmids in clinical settings[22]. The dynamics of these sweeps are governed by three mechanisms, each with its own contribution to plasmid fitness: horizontal transfer, within-cell competition, and between-cell competition. Perhaps the most well studied of these components, horizontal transfer, in which different bacteria exchange plasmids (Fig. 1A), porting diverse genes into new contexts[23], opening new evolutionary paths for their bacterial hosts[24–26]. However, beyond the sporadic action of horizontal transfer, plasmid evolution is continuously shaped by more fundamental forces acting at two scales. Between-cell competition is driven by differential replication of plasmid-host cells and within-cell competition is driven by replication interference between plasmids co-occurring in the same host cell, a phenomenon that often manifests as plasmid incompatibility[27] (Fig. 1B-C). Disentangling these two components of plasmid fitness is a central challenge to understanding plasmid ecology and evolution.

**Figure 1:**
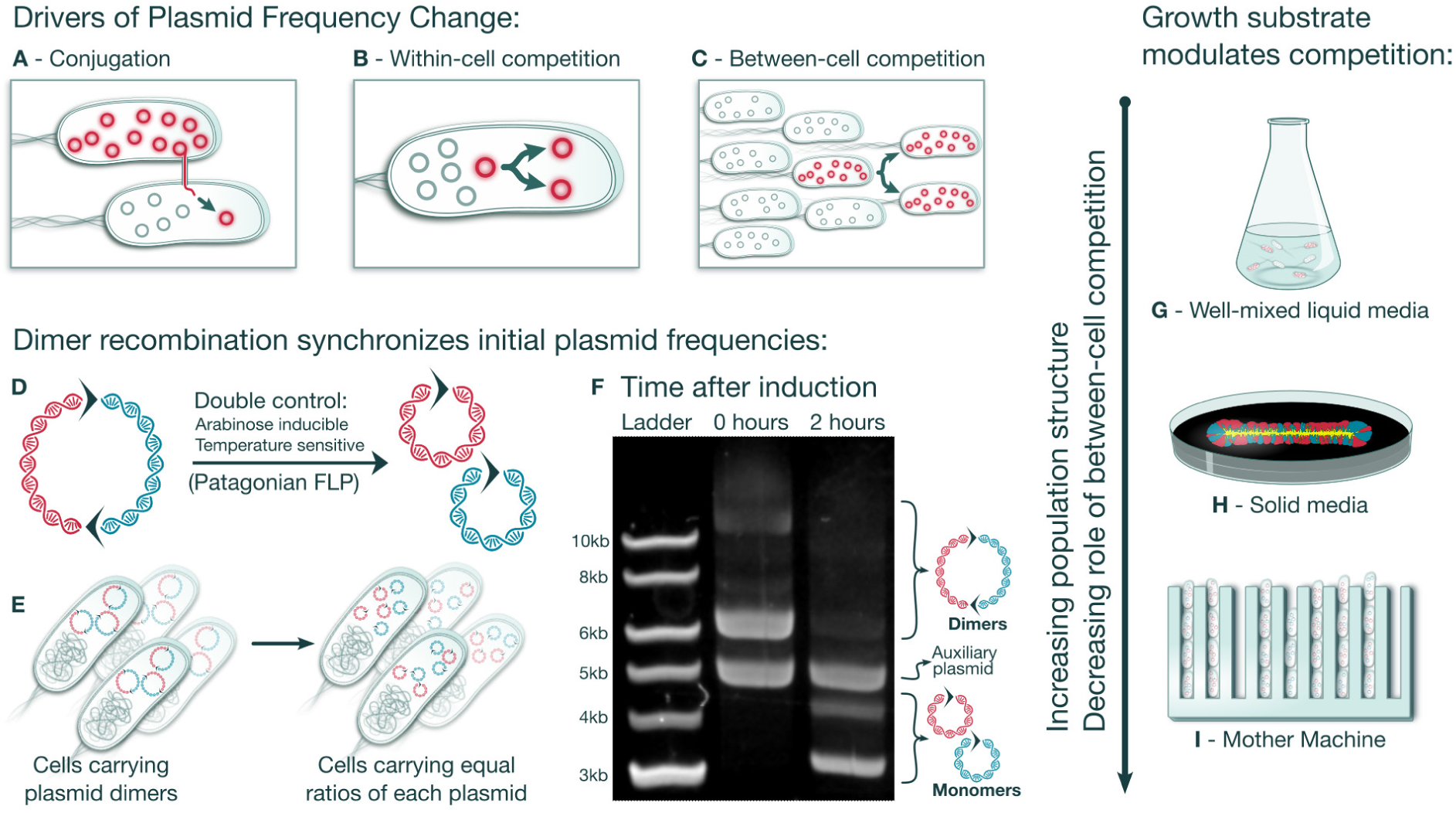
Synthetic plasmid dimers enable controlled within-cell competition experiments **A-C**, Schematic representation of plasmid fitness components. **D**, Illustration of synthetic plasmid dimers joined at collinear FRT sites represented by arrows. Activation of Patagonian FLP converts dimers into monomers, each containing one FRT site. **E**, Upon plasmid monomerization, bacterial cells carry equal frequencies of each competing plasmid. **F**, Plasmids extracted before FLP induction and after the addition of 0.3% arabinose and temperature reduction to 30°C were separated in a 0.7% Agarose-LAB gel. There is no visible plasmid monomerization before induction and efficient monomerization after induction. **G-I**, Schematic showing different cell culture modes with increasing spatial structure.

The contribution of within-cell processes to plasmid population dynamics is limited to the duration of the intracellular co-occurrence of competing plasmids. This co-occurrence is short-lived because cell divisions and random assortment of plasmids into daughter cells ultimately lead to each cell stochastically fixing one of the competing plasmids[27, 28]. Despite this constrained window of action, within-cell processes can play a critical role in plasmid evolution. This is because whenever a novel plasmid variant arises—by point mutation, gene acquisition, or cointegration with other plasmids, for example—it co-occurs intracellularly with the original variant[29], providing an opportunity for within-cell competition to act[5] even before the cell-level fitness effects become apparent.

Indeed, mounting evidence indicates that this transient plasmid co-occurrence is determinant to the frequency of plasmid evolutionary sweeps. For instance, higher plasmid copy numbers—which imply stricter within-cell competition—were shown to lower the fixation probabilities of novel plasmid variants[30] all the while slowing the rate of plasmid evolution[31]. Others demonstrated that a low dominance of plasmid encoded traits—which is only evident during plasmid co-occurrence—might also hamper plasmid evolution[4]. Additionally, plasmids with stable replicons were shown to be capable of displacing plasmids with unstable replicons[5]. Altogether these observations can only be explained by considering within-cell plasmid competition. Yet, the dynamics and temporal scales leading from the emergence of a novel plasmid to its within-cell fixation remain uncharacterized, despite their wideranging evolutionary, clinical, and economic importance.

These unknowns persist due to the technical hurdles in isolating and measuring the within-cell component of plasmid fitness. In general, previous experimental setups do not allow for the observation of changes in plasmid frequency immediately after the introduction of a novel plasmid variant[4, 5, 28, 30] or cannot quantitatively differentiate between plasmid frequency changes caused by between or within-cell competition[28]. As a result, major questions about plasmid population dynamics are still outstanding: First, we do not know how powerful within-cell genetic drift is (i.e. how fast do cells fix one of two equivalent competing plasmids). Second, we do not know which features of a plasmid might favor the within-cell fixation of one plasmid variant over another, nor do we know how deterministic the fixation of the favored variant is. Third, we do not know how within-cell genetic drift and intrinsic plasmid replication differences might interact with cell-level phenotypes (for instance, to which extent can a within-cell replication disadvantage of an emergent plasmid variant be offset by an increased cell-level fitness).

Here we address these questions by using a novel system for measuring intracellular plasmid competition dynamics based on synthetic dimerization and splitting of competing components. We first transform cells with two plasmid variants joined together in dimer form. Initially, the dimerization constraint couples the replication of both plasmid variants, ensuring that each variant is equally represented within each cell in the population. We then activate a recombinase that quickly converts dimers into monomers, uncoupling the replication of the plasmid variants and generating a population of cells each carrying independent plasmids of both variants in an equal proportion (Fig. 1D-F). This allows within-cell competition for replication to take place until the plasmid variants ultimately segregate. Note that, throughout this work, we use the concept of segregation in the Genetics sense—the appearance of cells carrying a single plasmid type (homoplasmic cells) from cells carrying both plasmid types (heteroplasmic cells) due to random copy allocation at division—rather than in the Plasmid Biology sense—the appearance of plasmid-free cells due to random partitioning. Additionally, we grow these cells in mother machines—microfluidic devices that minimize the population-dynamical effects of fitness differences between cells—allowing us to isolate the effects of within-cell plasmid fitness (Fig. 1G-I). By combining these techniques, we gain access to the transient plasmid population dynamics that were otherwise invisible in previous investigations of plasmid evolution.

## Results

### A system to measure within-cell plasmid competition by splitting synthetic dimers

We first sought to show that we could use the synthetic dimer method to create a population of cells each carrying independent, monomeric plasmids of two variants in equal proportion. We used restriction assembly to create plasmid heterodimers composed of two barcoded plasmids with PBR322 origins separated by collinear FRT sites (fig. S1A). We transformed those plasmids into DH5*α* cells also carrying an auxiliary, compatible plasmid encoding an arabinose inducible FLP recombinase[32]. The action of the FLP recombinase on the collinear FRT sites converted dimers into monomers (Fig. 1D), as is the case for similar, naturally occurring plasmid dimer resolution systems[33]. However, initial experimental pilots showed that leakiness of FLP expression was leading to premature dimer breakdown. To avoid this, we synthesized an *E. coli* codon-optimized version of the *Saccharomyces eubayanus* FLP gene.

*S. eubayanus* is a cryotolerant, Patagonian relative of the baker’s yeast *S. cerevisae*[34]. We hypothesized that, due to the perennially cold temperatures of Patagonia, the FLP enzyme of *S. eubayanus* might be thermosensitive, which we could use as an extra layer of activity control. We found that the *S. eubayanus* FLP was fully active at 30°C but completely inactive at 37°C. With this FLP variant, we were able to quickly convert plasmid dimers into monomers, but virtually no monomerization occurred before FLP induction (Fig. 1F). By sequencing barcodes across the FRT junctions we saw that about 96% of plasmids were in the monomeric form just 1.5 hours after arabinose induction. Moreover, the barcodes revealed that both plasmids were present in equal proportions across the bacterial population (fig. S1B,C), as expected. In all, we obtained a population of cells carrying a synchronized and equilibrated plasmid content, in which we could precisely measure subsequent plasmid content fluctuations, revealing the action of within-cell competition for replication.

### Quasi-neutral plasmid dynamics last for many days

We then set out to investigate the dynamics of quasi-neutral plasmid segregation and therefore characterize the typical duration of within-cell plasmid competition. In order to do so, we transformed DH5*α*+FLP with a heterodimeric plasmid composed of two monomers, pScar and pWater, identical but for a gene encoded fluorescent tag—mScarlet-I[35] and mWatermelon[36], chosen for differing at just 12 base pairs, having good spectral separation, strong brightness, and similarly fast maturation—and a restriction site used for dimer assembly. We used a razor blade to deposit a linear inoculum of dimer-carrying bacteria on the surface of solid LB+Ara (FLP-inducing) agar plates and let the bacteria grow at 37°C for 12 hours forming a linear expansion front, at which point the temperature was reduced to 30°C leading to the activation of the temperature-sensitive FLP and the monomerization of plasmids across the bacterial population. The expansion of the bacterial front was then monitored by time-lapse fluorescent imaging for at least 8 days (Fig. 2A), movie S1).

**Figure 2:**
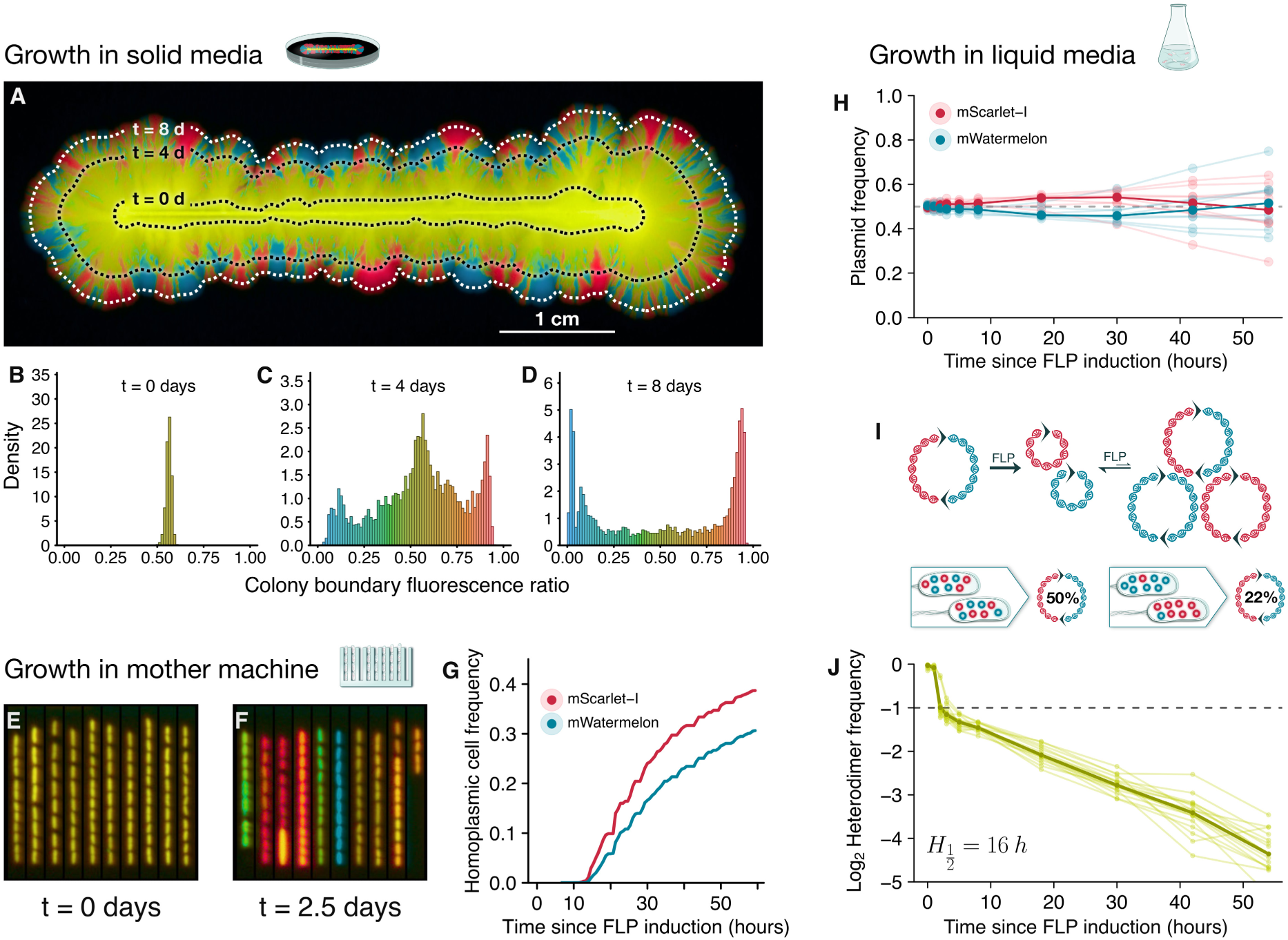
Quasi-neutral plasmid dynamics. **A**, Fluorescent image showing plasmid segregation as cells initially carrying equal proportions of pScar and pWater expand from the inoculation line for 9 days. Dotted lines show how far cells had expanded at each time point. **B**-**D**, Histograms of relative fluorescence across the colony boundary show that, at the time of FLP induction (**B**), cells fluoresced homogeneously in both colors, indicating equal carriage of competing plasmids. Further histograms (**C**,**D**) show the progressive emergence of homoplasmic sectors. **E**,**F**, Fluorescent images of pScar-pWater cells growing in a mother machine at the time of FLP induction (**E**), and 2.5 days later (**F**), highlight stochastic plasmid fixation. **G**, fraction of cells in mother machine that fixed each plasmid monomer type as a function of time. **H**-**J**, Cells carrying pScar-pWater dimers were grown in well mixed liquid media under FLP induction. Barcode sequencing at different time points revealed an initial increase in pScar frequencies that is consistent with a within-cell advantage to pScar plasmids (**H**). FLP induced, transient reformation of dimers can be used to assess average population heteroplasmy (**I**). Tracking heteroplasmy from the moment of FLP indution onwards reveals an initial decrease associated to within cell recombination equilibrium and a suqsequent exponential decay with *H* ½ = 16 hours or 22 cell divisions (**J**).

The time-lapses revealed a complex genetic drift pattern throughout bacterial expansion emerging from the interplay between plasmid segregation and demographic stochasticity. Immediately after FLP activation every cell at the expansion boundary contained equal proportions of each of the two competing plasmids, as indicated by the homogeneous fluorescence ratio across the entire boundary (Fig2B). However, subsequent cell divisions with random plasmid allocation produced daughter cells with varied plasmid compositions, some carrying a single plasmid type (homoplasmic cells), others still carrying both (heteroplasmic cells). Furthermore, demographic stochasticity at the expansion boundary led to the separation of cells into homoplasmic and heteroplasmic sectors (Fig. 2C). Finally, the remaining heteroplasmic cells continued to give rise to new homoplasmic cells and sectors, which ultimately overtook the boundary (Fig. 2D). Strikingly, this nested sectoring pattern is a signature of the double scale of plasmid genetic drift: when we inoculated an equal mix of both homoplasmic cell types instead of the dimer-carrying cells—effectively removing within-cell genetic drift and isolating the effect of demographic stochasticity—the bacterial expansion front almost immediately separated into coarse, homogeneous sectors (fig. S2), recapitulating prior studies[37]. The demixing of plasmids within the same -cell is much slower than the spatial demixing between cells as the colony expands and, thus, cell-level genetic drift contributes to the observed within-cell genetic drift.

To characterize the intrinsic rate of within-cell genetic drift in the absence of demographic stochasticity, we sought to experimentally isolate the within-cell plasmid dynamics from the effects of cell-level population dynamics. We populated the trenches of a mother machine, a microfluidic device for isolation of single-cell bacterial lineages[38], with dimer carrying cells and then induced plasmid monomerization (Fig. 2E). A key aspect of the mother machine is that, because mother cells are immobilized at the base of the trenches and cannot be displaced, every mother cell represents an independent lineage[39, 40]. At each cell division, plasmids are stochastically allocated between a daughter cell and a mother cell which remains at the base of the trench. This plasmid allocation is independent of any cell-level fitness effects, which therefore do not influence the probability of any of the two competing plasmids fixing in the mother cell. However, if one of the two plasmids has a replication advantage over the other, then that plasmid is more likely to increase in frequency and therefore fix in the mother cell. Ultimately, the mother cell fixation probability of each of the two competing plasmids is a measure of within-cell competition that is independent of between-cell competition. Unexpectedly, even though the competing plasmids were by design almost identical, pScar displayed a detectable within-cell competitive advantage over the pWater (Fig. 2F-G).

To exclude potential bias from fluorescence measurements, we developed an orthogonal method to quantify within-cell dynamics using DNA barcodes. Although mScarlet-I and mWatermelon have similar brightness and maturation profiles, the different spectral properties of the fluorophores could bias the detection of homoplasmic cells of each type, which would be erroneously interpreted as a within-cell fitness advantage to one of the plasmids. To rule out this possibility, we marked each plasmid variant with 10 independent DNA barcodes. We grew barcoded dimer-carrying cells in well-mixed, liquid LB medium and then induced plasmid monomerization. We kept liquid cultures at mid-exponential phase and collected samples at increasing time intervals from which we extracted the plasmids. Illumina sequencing of the barcodes revealed that immediately after monomerization, both plasmid variants were present in equal frequencies, after which pScar increased in frequency (Fig. 2H), corroborating its within-cell competitive advantage over pWater using a fluorescence independent method. Moreover, after the initial pScar increase, plasmid frequencies stabilized indicating that as plasmid segregation gave rise to homoplasmic cells, within-cell competition between the plasmid variants ceased as expected.

To measure the rate of plasmid segregation (i.e. the heteroplasmy decay rate *H* ½), we exploited the reversibility of the FLP mediated plasmid monomerization reaction. Although plasmid monomerization is entropically favored with about 95% of plasmids being in monomeric form after FLP induction, transient re-dimerization also occurs. Notably, while all the pre-induction dimers are pScar–pWater heterodimers, re-dimerization gives rise to three potential dimer types: the heterodimer and the homodimers pScar–pScar and pWater–pWater (Fig. 2I). We measured the proportion of heterodimers within total dimers for plasmid samples collected at each time point by electrophoretically separating the reformed dimers from monomers and sequencing the barcodes across the FRT junctions (fig. S1D,E). This heterodimer proportion (*hp*) encodes information about the average degree of plasmid segregation across the cell population. For instance, in the case of cells carrying both plasmid variants in equal frequencies, the expected probability that re-dimerization gives rise to a heterodimer should be, by Hardy-Weinberg Equilibrium (HWE), equal to 0.5. In contrast, cells with unequal plasmid frequencies should generate a smaller heterodimer proportion, and at the extreme, homoplasmic cells should generate no heterodimers (Fig. 2I).

Changes to *hp* occurred in two phases (Fig. 2J). Initially, in accordance with our synthetic initial condition, *hp* was approximately 1. Following induction, FLP action led to quick plasmid monomerization and re-dimerization causing *hp* to precipitously fall to 0.5, closely matching the HWE prediction. Importantly, the fast FLP action combined with HWE-matching mean *hp* is an accurate reporter of the average population heteroplasmy. Second, rounds of cell division and stochastic plasmid allocation progressively gave rise to increasingly segregated cells causing an exponential decay of *hp* (Fig. 2J). This allowed us to infer that *H* ½ is about 16 hours or 22 cell divisions under our experimental conditions, matching our observations that heteroplasmic cells persist for many days. This extended period of within-cell plasmid co-occurrence is on the same time-scale as many experimentally observed evolutionary sweeps[41], suggesting that during plasmid evolution there is ample time for between- and within-cell selection to interact and jointly contribute to evolutionary outcomes.

### Hemimethylation at the plasmid origin modulates within-cell drift and competition

With a quantitative description of within-cell plasmid genetic drift at hand, we decided to investigate if a simple model of cell division and plasmid replication could recapitulate the observed dynamics. We initially adopted a simplified mechanistic approach following [42]. We assumed that plasmids initiated replication at a constant rate *k_in_* throughout the cell cycle, but that these attempts were aborted with probability 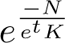 which arises from a multi-step inhibition process where is the total instantaneous plasmid density in a cell, and *K* represents the action of an inhibitor that is present in a concentration proportional to plasmid density. Note that when tracking two plasmid types within the same cell (say R and G plasmids), the stochastic replication of an R plasmid increases the subsequent total replication initiation rate of R plasmids because there are now more of those templates, all the while reducing the probability of successful replication of either plasmid types because of an increased action of inhibitor *K*. Altogether these two effects reduce the likelihood that the next replication event involves a G plasmid, effectively following a Pòlya urn process in which initial stochastic advantages are amplified during subsequent replications, leading to an increased variance and loss rate of co-occurring plasmids, even when the total density of plasmids remains stable within a cell. For simplicity, cells were initially assumed to divide after a constant time and upon division plasmids were binomially allocated to daughter cells. Although this is a two-parameter model (*k_in_* and *K*), because we are constrained by the empirical mean plasmid copy number at cell division (*N*^-^), the values of *k_in_* fully determine not only *K*, but also the variance *σ* of the stationary plasmid distribution and the autocorrelation *ϕ* between the plasmid copy number of a mother and a daughter cell (fig. S3A,B). When we tracked plasmid segregation in our simulations, we found that the average heteroplasmy of the cell population fell exponentially, qualitatively mirroring our experimental results (fig. S3C,D, movies S3-S4).

However, the model presented a quantitative discrepancy: simulated plasmid segregation occurred consistently faster than empirically observed, as evidenced by the theoretical values of *H* ½ obtained for the full range of *k_in_* covered (fig. S3E). We reasoned that by investigating model modifications that solve this discrepancy, we could point towards key processes that regulate plasmid dynamics in nature. First, we explored the role of mechanisms that reduce the randomness of plasmid partitioning (i.e. chromosomal volume exclusion, partitioning systems) by implementing models in which both daughter cells received exactly half of the mother cell’s total plasmids. Although this equal partitioning increased *H* ½, the magnitude of the effect was insufficient to reach the empirical value (fig. S3). Second, we observed that this simplified, effectively single-parameter model does not allow for the independent modulation of *σ* and *ϕ* (fig. S3F). Yet, population genetics indicates that the interplay between these parameters should contribute to the effective rate of genetic drift. Moreover, variability and autocorrelations in many cellular processes that our model ignores (i.e. interdivision times and expression states) might independently affect *σ* and *ϕ*. To investigate whether these effects might lead to a plausible *H* ½ we created a phenomenological model that allowed us to tune both *σ* and *ϕ* as independent parameters (see methods). Despite the flexibility of this model, all combinations in a biologically meaningful range of parameters led to overly fast *H* ½ (fig. S5A,S4A,B). Third, we turned our attention to a key assumption of the Pòlya urn replication dynamics: the possibility of immediate repeated plasmid replication. In reality, plasmids that just underwent replication might be unavailable for a subsequent replication round for some eclipsing period[42]. We incorporated this effect into our model by imposing that plasmids that recently replicated are less likely to be chosen to replicate again by a factor *ecl* ∈ [0, 1). We found out that a range of *ecl* values generated *H* ½ values consistent with experiments (fig. S5B,S4C), suggesting that eclipsing might be a key modulator of plasmid segregation, and therefore the inclusion of a system that delays repeated plasmid replication resolves the discrepancy between the model and observations.

Experimental ablation of eclipsing accelerated within-cell competition dynamics as predicted by our model. It is thought that eclipsing of ColE1-like plasmids might occur via hemimethylation driven sequestration[43, 44]. These plasmids have a conserved pair of DAM methylation sites near the origin of replication. When replication occurs, a fully methylated plasmid gives rise to a pair of hemimethylated plasmids. The origin-proximal hemimethylated DAM site pairs are then recognized and bound by seqA inhibiting further replications until full methylation is restored and the plasmids are released[45]. To ablate eclipsing we created pScarM^−^ and pWaterM^−^ by introducing a point mutation into each of the origin-proximal methylation sites.

Competition of pM^−^ plasmids in the mother machine revealed that plasmid segregation happened much faster than in the case of pM^+^ plasmid competition (fig. S5C). Naively, this finding might be counterintuitive: pM^−^ and pM^+^ plasmids achieve similar copy numbers[43], which are determinant to the rate of genetic drift and plasmid segregation [46]. However, in line with our theoretical results, an acceleration of segregation is expected due to increased replication variance caused by the uneclipsed Pòlya urn dynamics of pM^−^ plasmids. Moreover, our simulations showed that if a plasmid variant had an intrinsic replication advantage over another, such as a higher replication initiation rate, then eclipsing should act to reduce the fixation probability advantage of the better replicator (fig. S4D). This was empirically confirmed by the increased fixation advantage that mScarlet-I plasmids displayed in the combinations featuring absent methylation sites (fig. S5C). Altogether, hemimethylation-mediated eclipsing increased the duration of within-cell plasmid coexistence, all the while reducing the potential within-cell competitive differences between co-occurring plasmids.

### Plasmid transcriptional activity can reduce within-cell plasmid fitness

With the knowledge that ColE1-like plasmids contain hemimethylation sites that attenuate within-cell competitive differences, we asked which features of a plasmid could incur in consistently detectable within-cell fitness deficits despite this attenuation. We hypothesized that transcription-replication tradeoffs might affect plasmid replication efficiency[47]: DNA that is actively transcribed accumulates supercoiling which can inhibit plasmid replication in several ways[48]. For instance, supercoiling can stall DNA replication forks, delaying the completion of plasmid replication; supercoiling can affect the transcriptional activity that drives the production of plasmid regulatory and priming RNAs, decreasing replication initiation rates; and supercoiling can interfere with the annealing of primer RNA to the plasmid origin, also changing replication initiation rates. To test whether plasmid transcriptional activity reduces within-cell fitness, we created a new plasmid variant pScarA by replacing the strong ProC constitutive promoter that drives mScarlet-I production for the weaker ProA promoter[49] (Fig. 3A).

**Figure 3:**
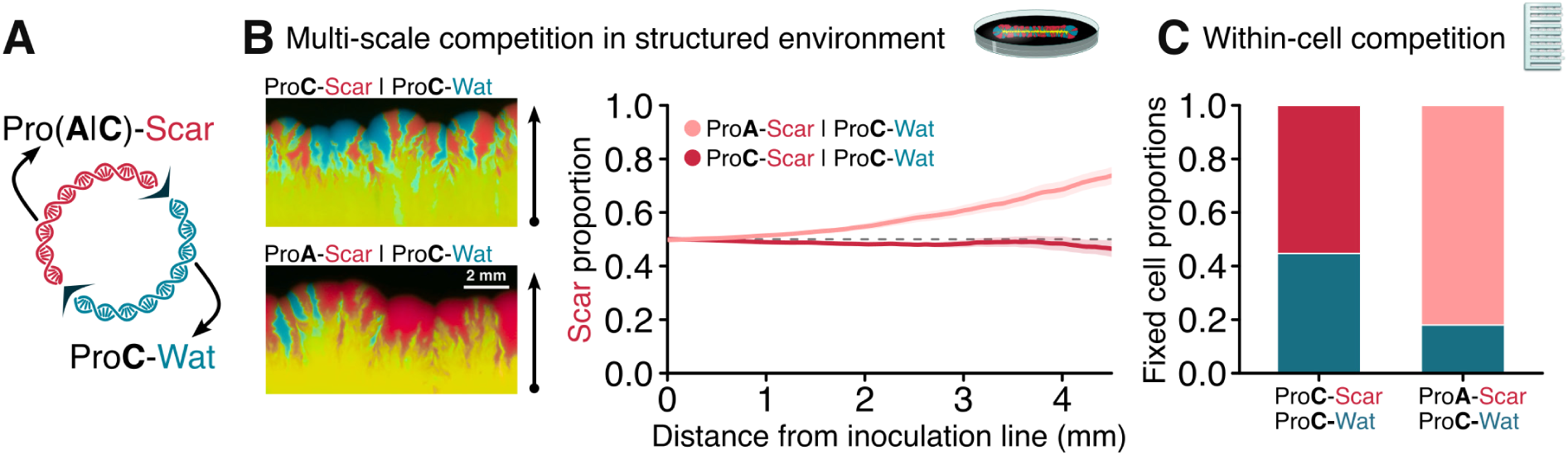
Transcriptional activity reduces within-cell fitness **A**, Schematic of a plasmid dimer composed of a monomer encoding mWatermelon under a strong promoter (ProC) and a monomer encoding mScarlet-I under either a strong or a weak (ProA) promoter. **B**, Fluorescent images show populations that expanded-under FLP induction-from a linear inoculum of cells initially carrying dimeric plasmids. When both monomers had fluorescent tags under the same promoter (pScar-pWater), the relative red fluorescence was constant throughout the population expansion, indicating neutrality. Instead, when one monomer had mScarlet-I under a weaker promoter (pScarA-pWater), the relative red fluorescence increased throughout the population expansion, indicating an advantage for the pScarA monomer. **C**, When dimer-carrying cells were grown in a mother machine under FLP induction till the fixation of one of the plasmid monomers, the less transcriptionally active pScarA was more likely to fix than the more transcriptionally active pScar, indicating that the observed advantage of pScarA is, at least partly, due to a within-cell competitive advantage.

Weakening transcriptional activity provided plasmids with a within-cell fitness advantage that allowed them to more easily displace their highly transcribed counterparts. In contrast to the relatively equilibrated outcome of pScar-pWater competition, when pScarA was competed against pWater using the dimerization method in solid agar medium, pScarA quickly overtook the more transcriptionally active pWater (Fig. 3B). It could be that this fast takeover was driven not by a within-cell replicative advantage of pScarA, but rather by the cell-level fitness cost incurred by higher fluorescent protein production from the highly transcribed pWater. While this cell-level fitness cost most certainly contributed to the final outcome of solid media experiments, we used mother machine experiments to isolate the contribution of within-cell competition to the expansion of pScarA. In the mother machine, the increased fixation probability of pScarA versus pWater (82%) when compared to the fixation probability of pScar versus pWater (55%) demonstrated that the reduction of promoter activity increased the within-cell fitness of pScarA (Fig. 3C). Overall, this revealed that plasmid gene expression—and all the cell-level phenotypic effects associated with that expression—might be in conflict with the within-cell fitness of the plasmid.

### The balance of withinand between-cell fitness depends on translational activity of plasmid mRNAs

This conflict between plasmid replication and gene expression implies that transcriptional activity might be costly for a plasmid on the within-cell scale of selection but beneficial on the between-cell scale. On the one hand, plasmid-encoded cell-level benefits (i.e. antibiotic resistance, heavy metal resistance, viral protection) are mediated by proteins that must be expressed from the plasmids, which leads to within-cell replication costs. On the other hand, these benefits allow plasmid bearing cells to outcompete plasmid free cells. To understand how these opposing levels of selection would guide plasmid population genetics, we constructed variants of pWater that encoded the trimethoprim (TMP) resistance gene dfrA following the mWatermelon fluorophore in a polycistronic operon driven by the strong promoter ProC.

The cell-level benefit conferred by the plasmid-borne dfrA gene is a product of the promoter activity—which incurs in a within-cell plasmid fitness cost—and the translational efficiency of the mRNA—which, in principle, should not directly affect plasmid replication. Indeed, we observed that the strength of the ribosomal binding site (RBS) governing dfrA translation affected cell-level fitness but had a negligible effect on within-cell plasmid fitness. We constructed two variants of dfrA encoding plasmids, one with a relatively weak RBS (pWater(w)dfrA) and one with a relatively strong RBS (pWater(s)dfrA)[50] which were then either transformed into cells by themselves of dimerized with the low transcription pScarA (Fig. 4A). Homoplasmic cells carrying pWater(w)dfrA, pWater(s)dfrA, or pScarA were grown together in liquid media with varying TMP concentrations. The growth rate of dfrA^+^ cells surpassed that of pScarA cells for higher TMP concentrations, with a consistently higher advantage for pWater(s)dfrA cells as expected from their increased dfrA translation (Fig. 4B). We then performed mother machine experiments with cells carrying pWater(w)dfrA-pScarA and pWater(s)dfrA-pScarA dimers at 0.08ug/mL of TMP—a concentration that strongly favored the growth of dfrA+ cells. In these experiments, mother cells fixed the low-transcription pScarA much more often than the cell fitness-enhancing dfrA^+^ plasmids (Fig. 4C), validating that the mother machine is indeed able to isolate the effects of between-cell fitness from within-cell fitness. Furthermore, the fixation rates of pWater(w)dfrA (10%) and pWater(s)dfrA (13%) were very similar, which is consistent with a within-cell fitness cost generated by their approximately equal transcription rate and independent of their translation-dependent cell-level benefits. In all, by manipulating plasmid transcription and translation rates, we were able to independently tune the betweenand withincell components of plasmid fitness.

**Figure 4:**
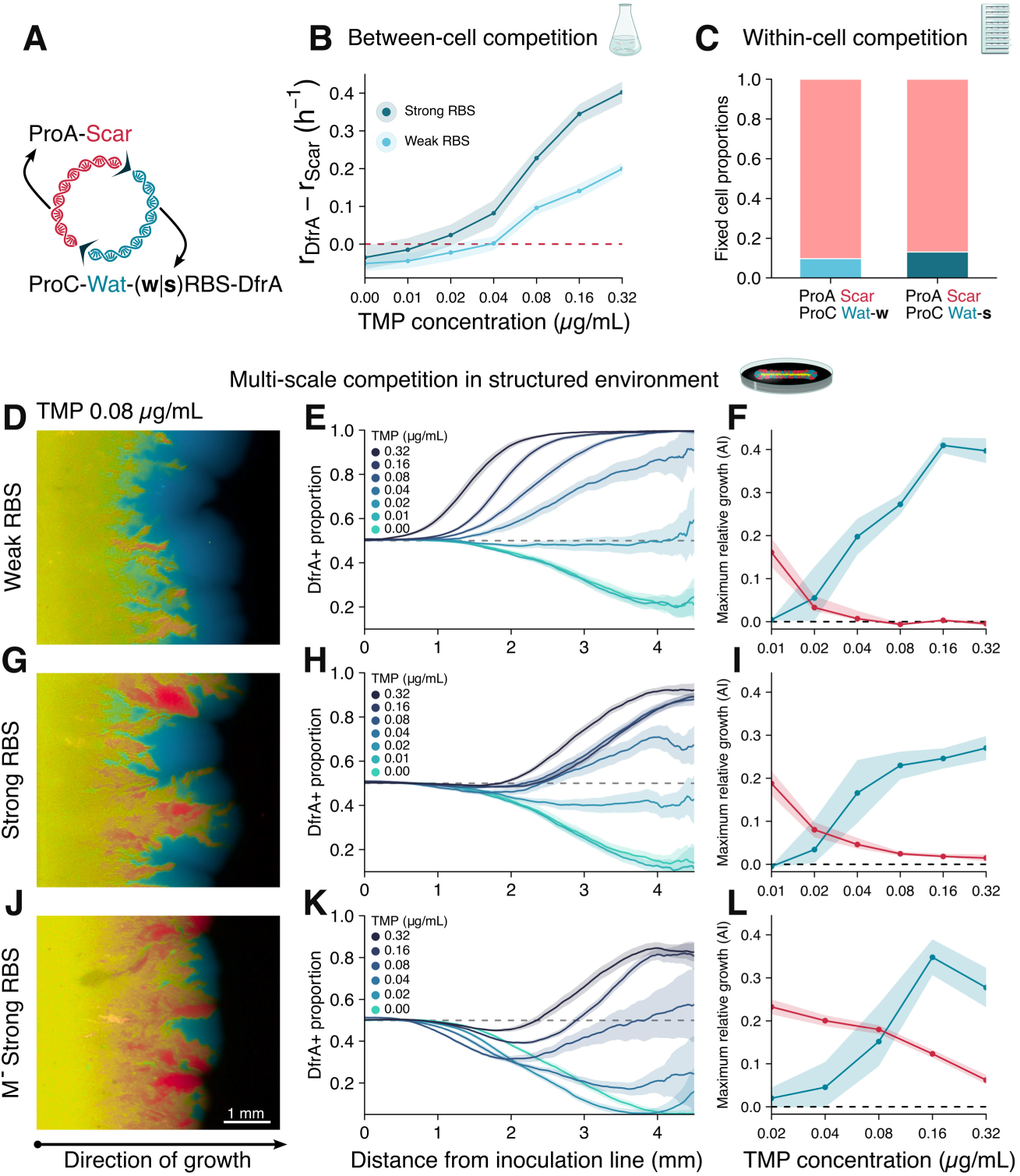
Within-cell competition may delay fixation of beneficial plasmid variants **A**, Schematic of a plasmid dimer composed of a monomer encoding mScarlet-I under a weak (ProA) promoter and a monomer encoding mWatermelon and the TMP resistence gene DfrA under a strong promoter (ProC). Two dimers were constructed, either with DfrA under a strong RBS or a weak RBS. **B**, Growth rates of cells carrying either pWater(s)dfrA or pWater(w)dfrA monomers relative to growth rates of pScarA-only cells in well mixed media with varying TMP concentrations. **C**, Plasmid fixation probabilities for cells initially carrying dimers and grown in a mother machine under 0.08µg/mL of TMP and FLP induction show that pScarA retains a within-cell advantage against pWater(w|s)dfrA, despite its fitness cost for the cell under those growth conditions. **D**-**L**, Populations expanding under FLP induction from a linear inoculum of cells initially carrying pScarA-pWater(w)dfrA (**D**-**F**), pScarA-pWater(s)dfrA (**GI**), or pScarA-pWaterM^−^(s)dfrA (**J**-**L**) dimers. **D**,**G**,**J**, Representative fluorescent images for cell grown under 0.08µg/mL of TMP. **E**,**H**,**K**, Normalized green fluorescence shows proportion of dfrA^+^ plasmids as a function of distance from inoculation line under several concentrations of TMP. **F**,**I**,**L**, Blue shows maximum increase in dfrA^+^ plasmid proportion in a spatial unit, and red shows maximum increase in pScarA plasmid proportion in a spatial unit. For TMP concentrations in which both the red and blue lines are bounded away from zero, a biphasic growth occurs, with an initial period dominated by a within-cell advantage for pScarA and a subsequent period dominated by the between-cell advantage for dfrA^+^ plasmids.

While experiments in well-mixed liquid medium remove spatial structure, which maximizes the relative importance of cell-level fitness[51], and mother machine experiments impose a rigid spatial structure, effectively nullifying cell-level fitness differences, in natural environments (i.e. enteric microvilli, biofilms) bacterial growth occurs with an intermediate degree of spatial structuring. To study the interplay of within- and between-cell levels of selection in the context of an intermediate degree of spatial structure, we performed pWater(w|s)dfrA-pScarA dimer competition experiments in solid agar medium with a range of TMP concentrations. When bacterial growth occurs in solid media, the reduced population size at the very boundary of the expanding bacterial colony increases the power of between-cell genetic drift, reducing the contributions of cell fitness to population dynamics compared to a well-mixed experiment[51, 52]. We found that, when starting from equal amounts of pScarA and either pWater(w)dfrA or pWater(s)dfrA within each cell, enough TMP selective pressure led to both of the dfrA^+^ plasmids overtaking the bacterial population (Fig. 4D-I). However, despite the common fixation outcome for dfrA^+^ plasmids, RBS strength profoundly altered the dynamics of plasmid fixation.

Although both dfrA^+^ plasmids displayed similar transcriptional within-cell fitness costs, the effects of within-cell selection were much more pronounced in the plasmid population dynamics of the higher expression pWater(s)dfrA than of the lower expression pWater(w)dfrA. In experiments performed within the range of TMP concentrations that favors dfrA^+^ cells, when pWater(w)dfrA was competed against pScarA, the proportion of dfrA^+^ plasmids (as measured by relative mScarlet-mWatermelon fluorescence) increased homogeneously as the bacterial population expanded from the inoculation line (Fig. 4D-F, fig. 4). This pattern is indicative of cell-level fitness being the major selection driver throughout the experiment. However, when the higher expression pWater(s)dfrA was competed against pScarA, fixation dynamics followed two distinctive phases. As the bacterial population first expanded, there was an overall increase in the proportion of pScarA plasmids. Yet, the stochastic nature of plasmid partitioning eventually gave rise to small pockets of cells dominated by dfrA^+^ plasmids. In the second phase, the emerging dfrA^+^ dominated sectors expanded against the pScarA sectors, driving an increase in the total proportion of dfrA^+^ plasmids (Fig. 4G-I, fig. 4). This biphasic pattern is consistent with an initial phase in which plasmid population dynamics are driven mostly by the within-cell plasmid fitness, causing an increase in the pScarA proportion across the cell population, followed by a secondary phase in which dynamics are driven by cell-level fitness, which promotes the expansion of dfrA^+^ sectors.

To confirm that the initial increase in pScarA proportion against pWater(s)dfrA was driven by a within-cell fitness advantage of pScarA, we conducted experiments that strengthened within-cell competition between those plasmids. We created pScarAM^−^ and pWater(s)dfrAM^−^ by ablating the eclipsing methylation sites. We expected that the stronger within-cell competition caused by Polyà urn plasmid replication dynamics of pM^−^ plasmids would lead to a longer, more pronounced initial phase of pScarAM^−^ proportion increase. As predicted, when the pScarAM^−^-pWater(s)dfrAM^−^ experiment was performed, the rate at which pScarAM^−^ proportions increased was higher and lasted for longer than in the pM^+^ competition experiments across all TMP concentrations (Fig. 4J-L). Although these experiments collectively demonstrated that within-cell fitness can drive plasmid population dynamics, they also revealed that these dynamics can not be trivially predicted by isolated measurements of within- and between-cell fitness: surprisingly, the competitive dynamics of pWater(s)dfrA were more hampered by within-cell competition than the dynamics of pWater(w)dfrA, despite pWater(s)dfrA conferring a higher cell-level fitness benefit at no extra within-cell fitness cost.

### The degree of trait genetic dominance mediates competitive effects at the withinand betweencell scales from plasmid invasion to fixation

To investigate the reason for the unexpectedly slower fixation of high cell-level benefit plasmids compared to low cell-level benefit plasmids in solid medium experiments, we built a spatially explicit model. Mimicking the experiments, cells were initialized at a central line in the simulated region. Cells at the boundary of the simulated colony were randomly chosen to replicate with a probability proportional to their relative fitness and their offspring cells were placed in an adjacent, cell-free position. Plasmids were partitioned between the daughter cells and then replicated as in our non-spatial model. We parametrized the relative fitness of homoplasmic cells using the data shown in Figure 4B and allowed pScarA plasmids to replicate with an intracellular advantage consistent with Figure 4C. As for the heteroplasmic cells, we assigned them relative fitnesses that were intermediates between those of the corresponding homoplasmic cell types. Crucially, we assumed that cell fitness could vary nonlinearly as a function of the proportion of dfrA^+^ plasmids it contained. This is because there are multiple levels of diminishing returns as the number of plasmid copies carrying a gene increases. For instance, ribosomes are limiting factors for protein synthesis, which is therefore sublinear in the mRNA concentration; once the TMP insensitive dihydrofolate synthetase encoded by dfrA is produced, the total enzymatic activity is also sublinear in the enzyme concentration; finally, the fitness benefits of the supplemental dihydrofolate produced by the synthetase are also sublinear. These conditions also imply that the fitness of cells carrying dfrA^+^ plasmids with a strong RBS must be more sublinear (more concave) than the fitness of cells carrying dfrA^+^ plasmids with a weak RBS (Fig. 4A-C). This is equivalent to saying that traits encoded by genes with stronger ribosomal binding sites should have a higher coefficient of dominance. Simulations indicated that the degree of trait genetic dominance was key to the rate of plasmid fixation.

We simulated equilibrated competition experiments between a plasmid that conferred no cell level fitness benefit—representing pScarA—and either a plasmid that conferred a high or a low fitness benefit—respectively representing pWater(s)dfrA and pWater(w)dfrA. When we assumed that heteroplasmic cell fitness was a linear function of plasmid composition, then the high-benefit plasmid always displaced the no-benefit plasmid faster than the low-benefit plasmid displaced the no-benefit plasmid (Sup fig), in contradiction to our experimental results. However, when the high-fitness plasmid was assigned a more concave fitness-response curve—as is expected from biological constraints—then its fixation rate dropped to below that of the low-fitness plasmid, matching the general aspect of the experimentally observed dynamics (Fig. 4B,E,H,K).

In our model, this dominance-dependent fixation rate of beneficial plasmids occured because the fixation is influenced not just by the fitness of homoplasmic cells (the endpoint fitness), but also by the fitness gradient (how much fitter is a cell that has a small increment of the beneficial plasmid proportion). The stronger dominance of high-benefit plasmids implies that small perturbations of plasmid content in cells at our initial condition—equal proportions of high- and no-benefit plasmids—cause negligible cell-level fitness differences (Fig. 4B), allowing within-cell fitness to dictate the initial dynamics (Fig. 4E). Conversely, the weaker dominance of low-benefit plasmids causes a steeper fitness gradient between cells differing by small perturbations in plasmid content (Fig. 4H). This steeper fitness gradient allows between-cell fitness to more efficiently counteract within-cell fitness, which leads to a faster fixation rate of low-benefit plasmids (Fig. 4K). Furthermore, our model also revealed that the role of dominance in the fixation of beneficial plasmids was dependent on the initial plasmid composition of the cell population.

When cells were initialized with a single copy of the beneficial plasmid against many copies of the no-benefit plasmid instead of the prior equilibrated competition simulations, the dominance-mediated slowdown of beneficial plasmid fixation was reversed. This initial condition is of particular biological interest because it represents the invasion by a novel plasmid variant that confers a cell-level benefit but is less replication-capable. Such an invasion could occur, for instance, if a plasmid captured a copy of a beneficial chromosomal gene, amplifying its effect and giving rise to a novel plasmid variant. The increased fixation of the strongly dominant high-benefit plasmid in invasion simulations can be explained by fitness gradients. When cells carried the high-benefit plasmid at a low proportion, the strong dominance caused a steep fitness gradient that ensured preferential replication of cells with additional copies of the high-benefit plasmid (Fig. 4A) and favored its fixation across the expanding cell population (Fig. 4D,J). Conversely, the less steep fitness gradient of low-benefit invading plasmids led to more loss and rarer fixation of the low-benefit plasmid (Fig. 4G,J). These theoretical results highlight a dual role for trait dominance in plasmid population genetics, favoring beneficial plasmid retention in the initial stages of an evolutionary sweep, but slowing down fixation in the later stages[53] (fig. S7).

Our simulations also revealed that cells containing the high-benefit plasmids might be more susceptible to invasion by no-benefit plasmids than cells containing low-benefit plasmids. In nature, such invasions might happen when a beneficial gene encoded in a plasmid is deleted, leading to the emergence of a novel, reduced plasmid variant. When we simulated this occurrence, the strong dominance of high-benefit plasmids allowed an invading no-benefit plasmid to initially increase in frequency without much fitness cost to the host cells (Fig. 4C). This increased frequency of the no-benefit plasmids then led to small pockets of cells that stochastically fixed the no-benefit plasmid, despite their cell-level relative cost (Fig. 4F,L). Conversely, when low-benefit plasmid containing cells were invaded by no-benefit plasmids, the fitness gradient caused by the presence of the no-benefit plasmid led to the quick loss of the no-benefit plasmid across the entire expanding cell population (Fig. 4I,L). Altogether, these invasion simulations revealed an apparently conflicting behavior: high-fitness plasmids should be both better than low-fitness plasmids plasmids at invading cells carrying no-benefit plasmids, but also more susceptible to be invaded by no-benefit plasmids.

To test these theoretical predictions about the mutual invasibility of plasmids, we developed a modified experimental setup that allowed for synchronous plasmid invasion experiments. While our synthetic dimers made it possible to characterize dynamics from an equilibrated initial condition—which is ideal for within-cell fitness measurements—they cannot generate an invasion-like initial condition. Instead, to achieve such an initial condition, we integrated either pWater(w)dfrA or pWater(s)dfrA flanked by FRT sites into the chromosome of *E. coli* already carrying pScarA (Fig. 4M). When FLP was induced, the integrated plasmids were excised from the chromosome as circularized, functional plasmids (Fig. 4N), at which point each cell in the population carried one or few dfrA+ plasmids—depending on the chromosome multiplicity at the time of excision—competing against multiple pScarA plasmids. Conversely, we also integrated pScarA into the chromosome of cells carrying either pWater(w)dfrA or pWater(s)dfrA, which upon FLP induction generated the reverse initial condition: rare pScarA competing against multiple dfrA^+^ plasmids. Therefore, the chromosomal integration of plasmids followed by FLP induced release allowed us to create a synthetic initial condition that mimics the initial stages of a natural plasmid invasion or evolutionary sweep.

Plasmid invasion experiments displayed close agreement with the theoretical predictions for the roles of trait dominance and within-cell fitness in beneficial plasmid fixation dynamics. The high-benefit pWater(s)dfrA was overall more efficient than the low benefit pWater(w)dfrA at invading cells carrying pScarA (Fig. 4O,Q,S, fig. 5), mirroring the reversal from the equilibrated competition results observed in our simulations. In both simulations and experiments, the steep fitness gradient of the invading high-benefit plasmids led to the fixation of the beneficial plasmid in sectors spread across the entire expanding cell population (Fig. 4O), whereas invading low-benefit plasmids were mostly defeated by within-cell selection, giving rise to sparse sectors of cells fixed for the beneficial plasmids which could only then expand (Fig. 4Q). Moreover, the no-benefit pScarA plasmid could transiently invade cells carrying the high-benefit pWater(s)dfrA but not the low-benefit pWater(w)dfrA (Fig. 4P,R,T, fig. 5). These results demonstrate the complex role of RBS strength in modulating within-cell competition throughout a plasmid selective sweep: a strong RBS can protect an emerging beneficial plasmid variant from being lost to within-cell competition during the initial phase of an evolutionary sweep; however, the strong RBS can also act to empower within-cell competition in later stages of the sweep, effectively delaying the fixation of the novel variant.

**Figure 5:**
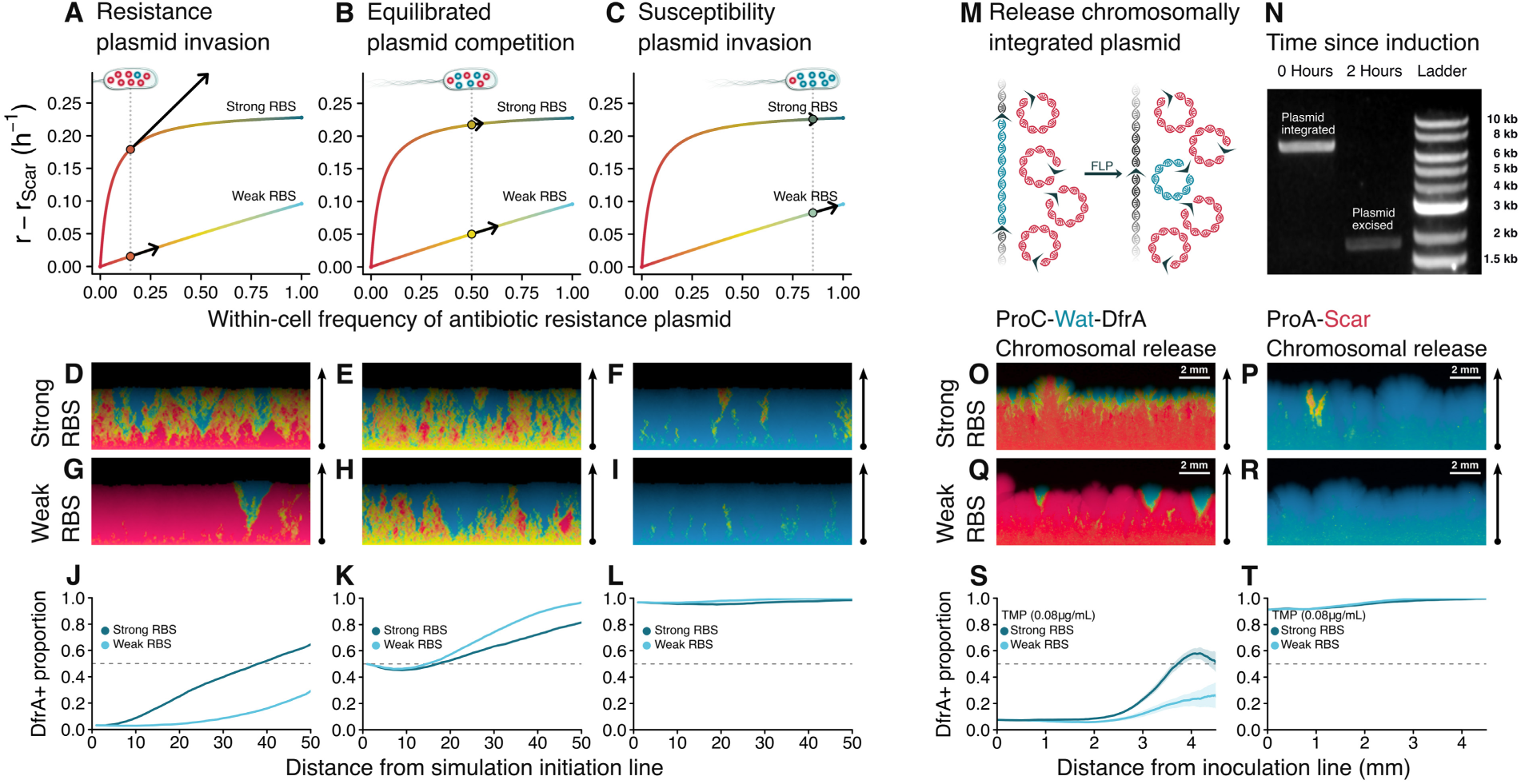
Spatial model of plasmid segregation predicts patterns of plasmid invasion **A**-**L**, Spatially explicit simulations of plasmid dynamics incorporating within-cell competition and between-cell competition were performed from three initial conditions: invasion of a beneficial plasmid representing the DfrA^+^ plasmids (**A**,**D**,**G**,**J**), equilibrated competition (**B**,**E**,**H**,**K**), and invasion of a costly plasmid (**C**,**F**,**I**,**L**). For invasion simulations, each cell was initialized with one copy of the invading plasmid competing against many of the resident plasmid. Two possible types of beneficial plasmids were considered: a high-benefit, high-dominance plasmid (representing a strong RBS), and a low-benefit, low-dominance plasmid (representing a weak RBS). **A**-**C** Show the relative fitness of cells as a function of their plasmid composition. Arrows show fitness gradients at those plasmid compositions. More dominant plasmids have higher fitness gradients when in low frequency, but low fitness gradients when in higher frequency. **D**-**I** Shows simulated bacterial expansion, with color representing their plasmid composition, blue being the beneficial plasmid and red the no-benefit plasmid. **J**-**L** Summarizes the simulations showing propotion of beneficial DfrA^+^ plasmids as a function of distance from simulation initiation line. **M**-**T**, Experiments with plasmid release from chromosome allow for empirical test of plasmid invasion. **M**, Schematic shows FLP induced release of chromosomally integrated plasmid to perform plasmid invasion experiment. **N**, Amplification of the locus of plasmid integration reveals efficient release of plasmid upon FLP induction. **O**-**R**, Fluorescent images show expansion from a linear inoculum (under FLP induction and 0.08µg/mL of TMP) of bacteria initially carrying a resident plasmid and a chromosomally integrated plasmid. **O**,**Q** Bacteria carry a resident pScarA and a chromosomally integrated pWater(w|s)dfrA. **P**,**R** Bacteria carry a resident pWater(w|s)dfrA and a chromosomally integrated pScarA. **S**,**T** Summarizes experiments by showing the relative fluorescence as a function of distance from inoculation line. As in the simulated invasions, the plasmids with stronger RBS were more likely to invade.

## Discussion

In this work, we developed a method that made it possible to measure the distinct withinand betweencell components of plasmid fitness. Two key issues had to be solved to conduct precise within-cell fitness measurements of competing plasmids: first, experiments required a controlled initial condition in which each cell contained equal proportions of the two competing plasmids and, second, fitness differences between cells carrying the different plasmids should not affect the measurement of within-cell fitness. We achieved this equilibrated initial condition by using a synthetic dimer system. Then, we used the geometric configuration of mother machine microfluidic devices to neutralize the effect of cell-level fitness differences. Although we initially investigated plasmids belonging to the widespread ColE1-like class, this method can be used on the broad group of theta-replicating plasmids, which includes—but isn’t limited to—IncF, RK2, ColE2-like, and PSC101-like plasmids. While some of the insights gained here might be specific to ColE1-like plasmid dynamics, the wide applicability of this approach will enable an in-depth characterization of the within- and between-cell evolutionary tradeoffs of many more plasmid classes.

A fundamental quantity underpinning the importance of within-cell competition for plasmid evolution is the duration of within-cell co-occurrence of competing plasmids. We determined that quasi-neutral plasmids can co-occur within the same cells for much longer than what was suggested by naive models of plasmid replication. The reason for this long coexistence was the presence of a hemimethylation based mechanism that was initially unaccounted for in our models. This mechanism functioned as a replication checkpoint that inhibited sequential replication events from a same plasmid, reducing the variance of plasmid replication and increasing co-occurrence duration. Incidentally, this mechanism also acted to reduce within-cell fitness differences between competing plasmids, which in turn made cell-level fitness relatively more important in determining plasmid dynamics.

The presence of conflict-mediating mechanisms that control fitness differences in one level of organization—in our case, the plasmid level—while ensuring that natural selection acts at a higher level of organization—in our case, the cell-level—is known as fitness alignment[54]. These mechanisms are often associated with systems undergoing the so called evolutionary transitions in complexity[1]—in which independent replicators coalesce into a coordinated replicating unit—and are ubiquitous in nature: for instance, queen bees sterilize their workers; eukaryotic chromosomes have synchronized replication initiation; multicellular organisms segregate somatic cells from germ cells[8]. The conservation of a hemimethylation-based replication control system across ColE1-like plasmids[43] is a sign of the evolutionary trend of fitness alignment between cell and plasmid[55, 56]. Crucially for plasmid evolution, these systems provide a reduction of within-cell fitness differences that promotes the retention of plasmids that would otherwise be relatively unfit within a cell, but might provide cell-level benefits.

Beyond the ColE1 class, these hemimethylation systems have been identified in highly divergent plasmids[57–60], pointing towards the potential broadness of this fitness alignment phenomenon. One reason for the convergent adoption of this hemimethylation mechanism across plasmid classes might be the ease with which few mutations can generate regulatory methylation sites: while the proteinaceous components of the system—such as the methylases and sequestration proteins—are often encoded in the bacterial genome, a plasmid only requires two adequately spaced, origin-proximal, four base-pair long methylation sites to display the replication eclipsing that underlies fitness alignment. Notably, the stringency—and consequently the fitness alignment effect—of these hemimethylation mechanisms varies substantially across plasmids. On one end of the spectrum, stringent plasmid hemimethylation mechanisms lead to replication coupling to the chromosome and tight copy number control[59, 60]—which is a strong form of fitness alignment[54]—while, on the other end of the spectrum, lax hemimethylation mechanisms simply delay repeated plasmid replication, potentially reducing the within-cell competitive effects without necessarily affecting copy number[43, 58]—a much more subtle form of fitness alignment.

Despite the existence of this within-cell competition regulatory system, we found that the transcriptional activity of a plasmid incurs in a within-cell fitness cost that might cause it to be displaced by a less transcriptionally active plasmid variant. This transcriptional burden leads to a fundamental tradeoff between plasmid-level and cell-level fitness: cell-level benefits conferred by plasmids are contingent on protein expression—for instance, of an antibiotic resistance gene such as dfrA—which in turn is costly for plasmid replication. Because protein expression is a product of transcription and translation but only transcription directly affects plasmid replication, we were able to independently tune the within- and between-cell components of plasmid fitness by manipulating promoter activity and RBS strength. In doing so we uncovered an unexpected evolutionary role for RBS strength that was mediated by the degree of plasmid trait dominance.

A stronger RBS driving dfrA translation caused a more dominant plasmid-encoded antibiotic resistance trait, which had opposing effects in the early stages versus in the late stages of a selective sweep. Initially, strong trait dominance favored the increase of cell-beneficial invading plasmids against a resident type that was a better within-cell competitor. However, in later stages of the selective sweep, stronger trait dominance empowered within-cell selection and prevented the fixation of the cell-beneficial plasmid. Although the role of trait dominance is well-established in classic population genetics, plasmids present a much more elaborate situation[4]: in diploid organisms, three numbers—the fitnesses of the heterozygote and of the two homozygotes—suffice to characterize the invasion of a genetic variant[53]; by contrast, for bacterial plasmid variants, an entire curve—which associates each potential plasmid composition to a fitness value—contributes to a selective sweep. This fitness profile further interacts with within-cell competition and stochasticity in plasmid replication and partitioning, jointly governing the complex dynamics of plasmid invasion. Taken together, our experimental and theoretical results suggest that an intermediate degree of dominance must promote a maximal plasmid invasion efficiency by balancing early and late stage invasion benefits. However, the exact shape of an optimal fitness curve remains an open problem and probably depends in a non-trivial manner on the bacterial population structure, the strength of within-cell competition, and the timescale considered.

Overall, we found that within-cell competition was able to effectively oppose the fixation of invading plasmids that would otherwise have been beneficial for cells, particularly in spatially structured situations with smaller cell populations and strong cell-level genetic drift. Plasmid evolution in natural environments must balance these conflicting demands at within- and between-cell scales. Because the relative importance of within-cell competition was modulated by promoter activity and RBS strength, we expect that transcriptional and translational activities from plasmid-bound genes must differ from those of chromosomal genes. Indeed, it is a well-known pattern that plasmids are more AT rich and less transcriptionally active than chromosomes. This has often been ascribed to cell-level fitness: the translational burden of a plasmid might reduce cell fitness, so both cells and plasmids coevolved to silence plasmid genes[61, 62]. However, our results suggest that the pattern might also be explained from the perspective of within-cell plasmid fitness: transcription is at odds with plasmid replication, so plasmids evolved higher AT content that allowed them to co-opt the bacterial H-NS gene-silencing machinery. Two lines of evidence lend credence to this plasmid-driven silencing hypothesis: first, previous studies have suggested that the fitness cost to cells upon plasmid acquisition is not driven by plasmid translational burden[63, 64], but rather by a large dysregulation of chromosomal gene expression[64]; second, plasmids oftentimes carry self-silencing H-NS paralogs[65], but this regulation always happens in the level of transcription and not translation.

Finally, our results raise new hypotheses to explain some of the mysteries surrounding the so-called cryptic plasmids. Recent metagenomic analyses have revealed that many of the most common replicons in human microbiomes are small, multicopy plasmids that are almost devoid of transcriptional activity and provide no obvious benefit to their host cells[13]. Our work suggests that the emergence of such minimal plasmids should be greatly accelerated by within-cell competition, particularly for multicopy replicons. Starting from an ancestral plasmid with functional genes, higher copy numbers should increase the rate at which plasmids with a gene deletion appear. After this initial appearance, higher copy numbers should make the results of within-cell dynamics less stochastic, favoring the fixation of the less transcriptionally active plasmid variant harboring the deletion. This within-cell competitive superiority of minimal plasmids might provide an indirect fitness benefit for their host cells. By outcompeting other plasmids that attempt to conjugate into their hosts, these minimal plasmids might function as a defense system against potentially costly horizontally transferred genes.

## Methods

### Strains and media

All experiments were performed in DH5*α* cells carrying an arabinose inducible Patagonian FLP and a chloramphenicol resistance cassette. All other plasmids also carried a kanamycin resistance cassette. All experiments were performed in LB and in the presence of 20µg/mL chloramphenicol and 50µg/mL kanamycin to prevent plasmid loss.

### Time series in well-mixed media

In LB medium, we grew a mix of bacteria with barcoded dimeric plasmids to OD_600_ = 0.5 at 37°C. Dimer splitting was induced by adding arabinose to a concentration of 0.3% and moving cell culture to 30°C. OD_600_ was kept below 1 at all times and whenever passaging, arabinose concentration was kept. To track plasmid composition, at every time point we minipreped 10mL of culture at OD_600_ *>* 0.4. Plasmid frequencies were then assessed by Illumina sequencing of barcodes. 10 replicates were performed with distinct barcode combinations. NEB Monarch kits were used for minipreps and DNA cleanup.

### Heteroplasmy quantification

To track segregation level it is necessary to increase the relative yield of dimeric plasmid versus monomeric plasmid. To do so we thermally inactivated the FLP recombinase and let cells grow. The intrinsic replication advantage of dimers (double origin) then increases the dimer yield. At every time point we took 1mL of culture and inoculated it into 50mL of LB. We grew this mix at 42°C for 2 hours. Then, in a 15mL conical, we mixed 6mL of culture with 4g of crushed ice to immediately chill culture and inhibit any FLP recombination activity. We proceeded to miniprep keeping chilled until the cell lysis step (cold centrifugation step and cold cell resuspension buffer). We resuspended the miniprep product in 40µL. To separate dimers, we ran 5µL of the miniprep product in a 0.7% agarose LAB gel for 45 minutes at 12.5V/cm. We gel extracted the band corresponding to the dimeric plasmid and ressuspended the DNA in 30µL. To “Invert” the barcodes, we digested 0.5ng/µL of dimeric plasmid with EcoRI and Mung Bean endonuclease in 20µL of CutSmart buffer for 3 hours at 30°C. Then we completed the volume up to 100µL with T4 ligase buffer and 4µL of T4 ligase and let blunt religation reaction proceed for 3 hours at 22°C. We cleaned DNA and resuspended in 6µL of nuclease free water. We used 1µL in a 25µL PCR to add Illumina adapters and timestep barcodes for 12 cycles.

### Solid media experiments

Solid media was prepared by autoclaving 2% Bacto agar, 2% LB, 1% Mars Black pigment, and then adding a filtered arabinose solution so that the final arabinose concentration was of 0.3%. 40mL were poured in each petri dish and an overlay of 5mL of 0.5% Bacto agar was added on top. When a TMP (in a DMSO solution) was added, care was taken such that every dish had the same concentration of DMSO regardless of TMP concentration. Inoculation of cells was performed by wetting a razor blade with a saturated cell culture, touching the blade twice on a dummy agar plate to remove excess liquid, and then touching the target plate. Inoculated plates were grown for 12 hours at 37°C (in which FLP is inactive) and then transferred to a moist incubator at 30°C for up to 2 weeks.

### Image acquisition and analysis

Image acquisition followed [66]. Red channels and green channels were aligned, background fluorescence was subtracted from the images, and each channel was re-normalized such that the median fluorescence of each channel in a region around the inoculation line was equal. This is because, in that central region, cell growth occurred prior to dimer splitting and, therefore, the fluorescence ratio in that region represents equal plasmid frequencies. Then, at each pixel *i*, the proportion of red (green) plasmid *p_R,i_* (*p_G,i_*) was calculated as 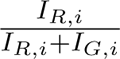 where *I_R,i_* is the re-normalized red intensity at pixel *i*, and *I_G,i_* is the re-normalized intensity at pixel *i*. The proportion of red plasmid as a function of the distance inoculation line was defined as 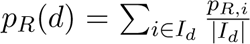 where *I_d_* is the set of pixels that are at a distance *d_d_* from the inoculation line, and |*I_d_*| is the number of such pixels.

Confidence intervals were built by a block resampling approach. For each experimental condition, images were randomly broken into 1cm blocks. This is because if cells are more than a centimeter apart, their dynamics are statistically independent: they do not descend from each other nor do they directly compete for resources in any way. From randomly resampling these blocks, synthetic data sets were constructed, and statistics recalculated. From these resampled statistics, confidence intervals were produced.

### Mother machine experiments

A mixture of 25*g*: 2.5*g* silicon-base:curing agent (sylgard-184-elastomer) was used to cast the pattern from the master wafer[67]. A 700*µm* diameter coring needle was used to punch out inlets and outlets and the device was cut out of the cast, with a razor blade. The device and coverslips were heated to 37°C and the surface was treated using *O*_2_ plasma before bringing them into contact, then baked at 95°C for 1 hour. The microfluidic chips were then stored at room temperature until use. Before loading the cells into the microfluidic the device was primed using ∼ 200*µL* of Biofloat Flex coating solution (faCellitate, F202005), and was left to rest for 30 minutes.

The dimer-carrying cells were grown to saturation, centrifuged at 5000 g for 30 seconds, and concentrated ∼ 10*X* by removing the supernatant and resuspending the pellet in ∼ 200*µL* of their respective cultures.

Two distinct cell mixes were loaded in each mother machine run. The first concentrated culture was then injected into the microfluidic and centrifuged into the top rows of the chip using custom adapters to spin the device on a table top centrifuge for 1 minute 650 g. Then the first culture was washed away, the second concentrated culture was loaded and using the same procedure it was seeded on the opposite trenches.

Then the device was connected to a media source that pumped it through the chip using a peristaltic pump at a rate of ∼ 260*µL/min* and into a waste container. Media was prepared by growing cells in LB up to OD_600_ = 0.5, spinning down and filtering out cells, and adding PEG-PPG-PEG to a final concentration of 0.05%. The use of partially spent media ensured that cells within the mother machine experienced a similar chemical environment than those growing in free bulk conditions.

The whole setup was transferred to a microscope and kept at 30°C for imaging. Every position was imaged every 40 minutes in the fluorescent channels corresponding to the proteins in the strains, and the total intensity of each trench containing one clonal micro-colony was extracted from the images. Cells were first grown for 4 hours, at which point each trench was populated by cells descending from a single mother cell. Then filtered arabinose was mixed to a final concentration of 0.3% promoting the splitting of the dimers. Cells were then imaged for a period of up to 4 days to track plasmid segregation.

### Mechanistic model

First, for each combination of *K* and *k_in_* the stationary plasmid copy number distribution (conditioned on the presence of plasmids) was computed by iteratively solving in continuous time the master equation *dP* (*N, t*)

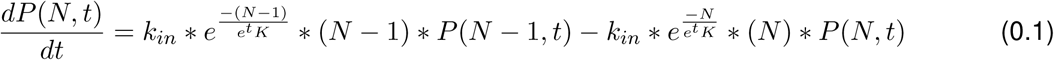

From *t* = 0 to *t* = 1, where *P* (*N, t*) is the proportion of cells carrying *N* plasmids at time *t*, and then applying a binomial redistribution kernel such that

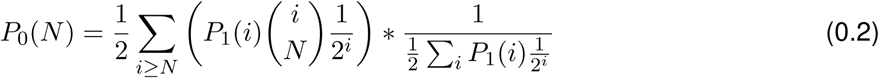

Where *P*_0_(*N*) is the initial proportion of cells carrying *N >* 0 plasmids at the beginning of the subsequent replication round and *P*_1_(*i*) is the final proportion of cells carrying *i* plasmids at the end of the last plasmid replication round. Once a stationary copy number distribution was found, an initial condition with equal copy numbers of R and G plasmids was built by setting

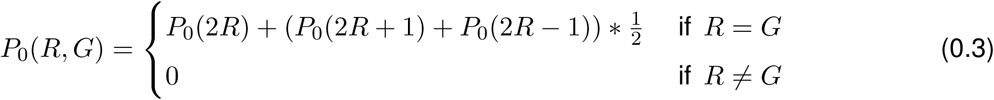

To solve for the time evolution of this 2D array of density function we do not need to solve a 2D array of Master equations. This is because the behavior of the master equation is independent of the identity of plasmids R and G. Instead, we can solve for the probability distribution *P^Ni^* (*N*) of ending a replication cycle with *N* plasmids from an initial cell carrying *N_i_* plasmids. The Δ*N* can then be allocated between R and G plasmids following a beta-binomial distribution, which is the solution of the Pólya urn process. In this case the density distribution of cell types after the replication cycle *P*_1_(*R, G*) can be given by

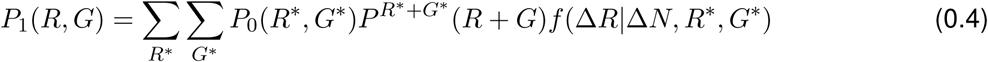

Where *P*_0_ (*R, G*) is the initial cell-type distribution, *P^R^*^∗+*G*∗^ (*R* + *G*) is the probability of ending with *R* + *G* plasmids conditional on starting with *R*^∗^ + *G*^∗^ plasmids (note that this quantity only depends on the sum and therefore does not have to be calculated a quadratic amount of times), and *f* (Δ*R*|Δ*N, R*^∗^*, G*^∗^) is the beta-binomial calculated at Δ*R* = *R* − *R*^∗^ with parameters Δ*N* = *R* + *G* − *R*^∗^ − *G*^∗^*, R*^∗^*, G*^∗^.

For the model with equal plasmid allocation between daughters at cell division, the binomial redistribution kernel was replaced by

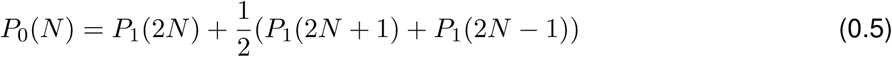

### Phenomenological simulations

In our model, plasmid copy number at cell division (PCNcd) was log-normally distributed across cells[68–70]. Throughout simulations we assumed the mean PCNcd was *N*^-^ = 40, a value consistent with the empirical average plasmid copy number (note that average copy number at cell division is a larger number than average copy number, since plasmid duplications occur throughout the cell cycle), and was distributed with a variance *σ*^2^ *>* 0 as a free parameter. As a second free parameter, we also allowed for an autocorrelation *ϕ* ∈ [0, 1] between the PCNcd of a mother and its daughter cell. In this context, *σ* encodes how variation in transcriptional activity, RNA degradation, cell size and division rates, amongst other factors lead to PCNcd variability whereas *ϕ* encodes how well-correlated are those factors across cell generations. We initialized cells with an equal number of plasmids of two potentially distinct types (R & G) and at every simulated cell division we binomially allocated plasmids into the two daughter cells. Then, plasmids were randomly chosen to replicate until each daughter cell reached a target PCNcd. At every plasmid replication event, the probabilities of a R plasmid (*ρ_R_*) or of a G plasmid being chosen (*ρ_G_*) were given by

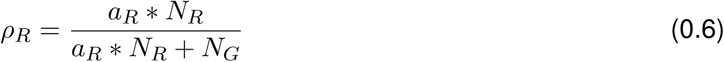

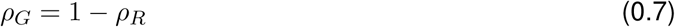

Where *N_R_* is the number of R plasmids in that cell, *N_G_* is the number of G plasmids in that cell, and *a_R_ >* 0 is a within-cell fitness differential between plasmids. When *a_R_* = 1 both plasmids have neutral within-cel dynamics, when *a_R_ >* 1 there is a within-cell advantage for R plasmids, and when *a_R_ <* 1 there is a within-cell advantage for G plasmids.

The target PCNcd was determined as to respect *N*^-^, *σ*, and *ϕ* as well as the log-normal stationary distribution of plasmid copy number. In other words the successive cell generations had randomly fluctuating plasmid numbers all the while respecting an autocorrelation with the copy number of the previous generation. Effectively, the PCNcd of a daughter cell (*N_t_*_+1_) was related to that of its mother cell (*N_t_*) by the equation

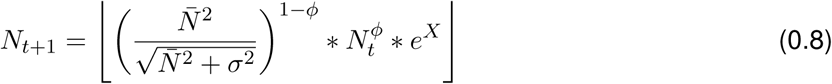

Where *X* is a normally distributed random variable following

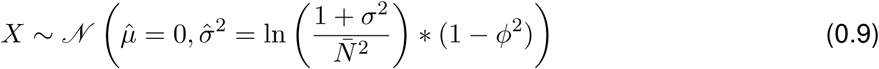

To incorporate plasmid eclipsing, we introduced a constant *ecl* ∈ [0, 1) imposing a probability penalty for the replication of plasmids that had already replicated in a cell cycle. The modified probability of a R plasmid being chosen to replicate was then given by

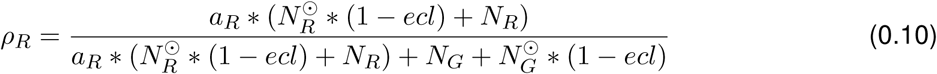

Where *N_R_* and *N_G_* now represent the numbers of plasmids that have not replicated in the ongoing cell cycle, whereas *N* ^⊙^ and *N* ^⊙^ represent the numbers of plasmids of each type that have already replicated at least once. In the case when there is no eclipsing (*ecl* = 0), plasmid replication follows a Polyà urn. When there is strong eclipsing (*ecl* → 1), then during a cell cycle each plasmid replicates at least once before any plasmid has a chance for a double replication.

For the heteroplasmy decay analysis, a population of *M* cells (4000 for most simulations) that divided at each time step was simulated. Heteroplasmy of the *i^th^* cell at the time point *t* was given by

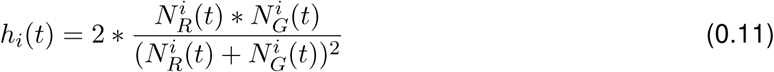

Where *N^i^* (*t*) (or *N^i^* (*t*)) is the number of R (or G) plasmids in the *i^th^* cell at the time point *t*. The population heteroplasmy was given by

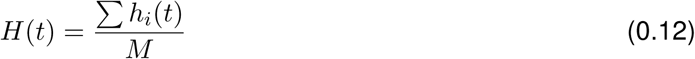

Note that under this definition, when all cells have equal amounts of each plasmid type, *H* = <u>^1^</u>, and when all cells are homoplasmic, *H* = 0. The asymptotic decay rate of heteroplasmy is given by

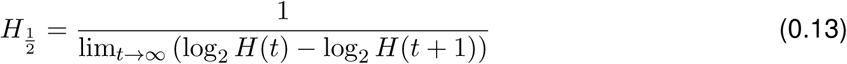

Which in practice we approximated by calculating *H* ½ at *t* = 50 generations.

For spatially explicit simulations, cells were initialized-with equal amounts of G and R plasmids-along a line in a square grid, mimicking razor blade inoculations. Plasmid replication dynamics followed the rules set in the non-spatial simulations. Without loss of generality we assumed that cells carrying only G plasmids were fitter (divided faster) than those carrying only R cells. In this formulation, G plasmids represent the dfrA^+^ plasmids for cells in the presence of TMP. Whenever a cell *i* was initialized with a frequency of G plasmids 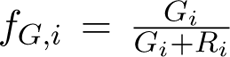, it was assigned a random, log-normally distributed time-todivision with standard deviation *σ*- = 0.2[71] and average value *T*^-^*_i_*, which was a function of it’s fitness advantage relative to an all-R cell Δ*^i^*:

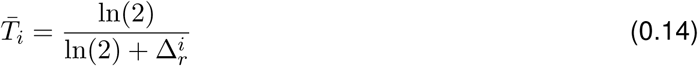

Δ*^i^* itself was a function of the cell plasmid composition:

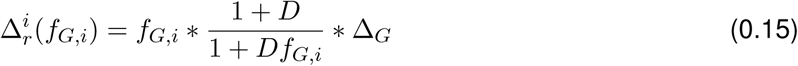

Where Δ*_G_*was the fitness advantage of an all-G cell relative to an all-R cell, and *D* is a measure of dominance of the plasmid encoded advantage. For *D* = 0 cell fitness linearly interpolates between all-R and all-G cells, and for increasing *d*, smaller proportions of G plasmids are needed for a fitness benefit. Values for Δ*_G_* were empirically determined.

## Supporting information

DNA Sequences of synthetic constructs

## Acknowledgements

We thank Noah Olsman, Akos Nyerges, Jana Huisman, Célia Souque, Siân Owen, Anurag Limdi, Sophia Wiesenfeld, Kepler Mears, and the rest of the Baym Lab for discussions and technical advice. This work was supported by the NIGMS of the National Institutes of Health (R35GM133700), the David and Lucile Packard Foundation, the Pew Charitable Trusts, Alfred P. Sloan Foundation and NSF grant MCB2426105.

## Supplementary Figures

**Figure S1:**
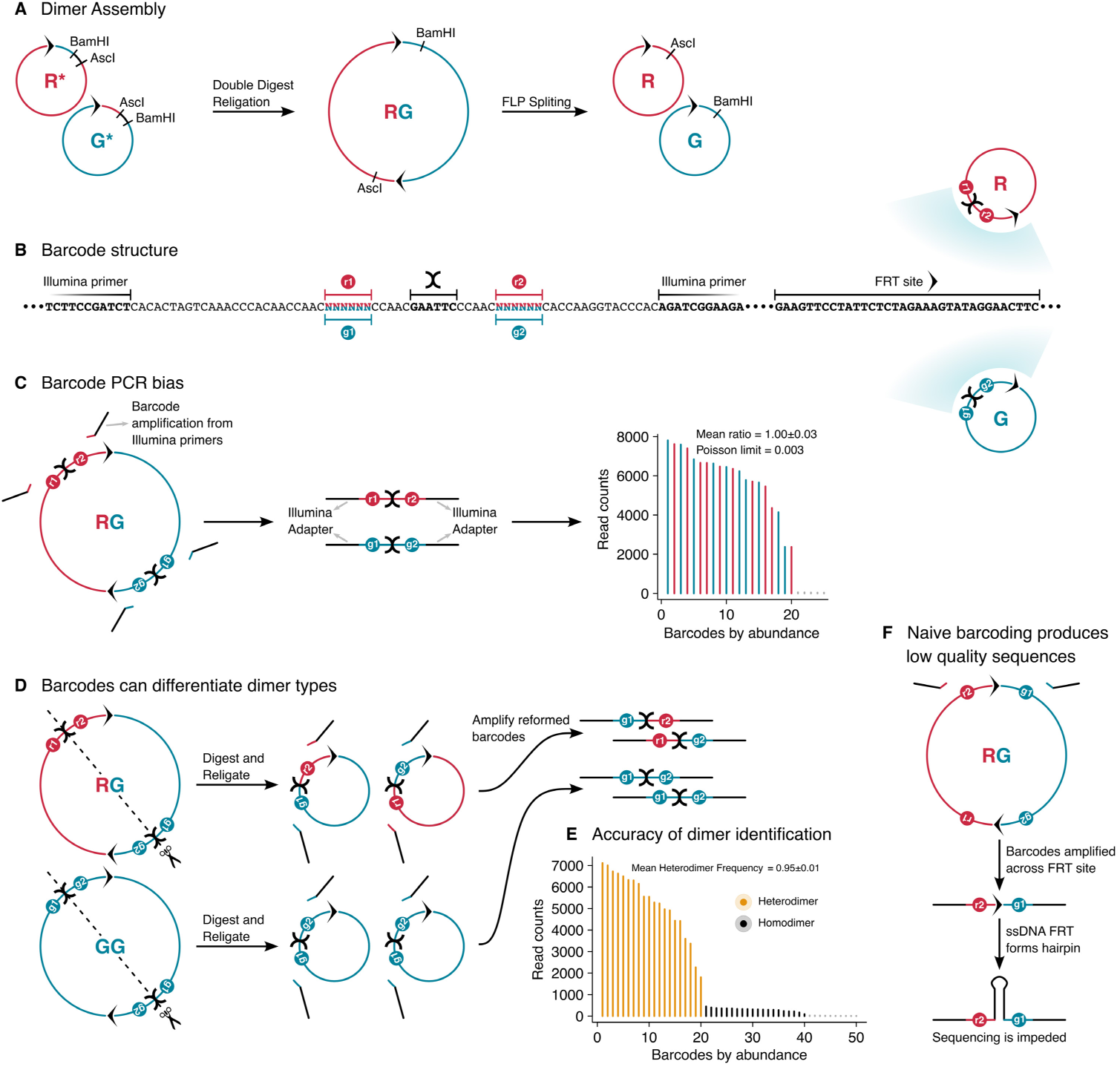
Dimer construction and barcoding **A**, Plasmid dimers RG were constructed from arbitrary monomers R and G by adding a pair of restriction sites (BamHI and AscI) at the same position but in reverse order for each monomer. A double digestion and religation creates the dimer, and upon FLP induction the monomers are regenerated within the cell, bearing a restriction scar. **B**, Plasmids were also barcoded. Each monomer received two barcodes separated by an EcoRI site and flanked by Illumina primer sequences that allow for simple downstream barcode sequencing. **C**, Dimers provide an internal control for PCR bias in barcode amplification. We mixed 10 independently barcoded dimers and amplified and sequenced barcode regions from that mix. Ratios between counts of barcodes belonging to the same dimer were on average 1.00 0.03. **D**, To discriminate heterodimers from homodimers, purified dimers were cut at the EcoRI site and religated, uniting barcodes from each side of the dimer. Those reformed barcodes were amplified and sequenced. **E**, Accuracy of dimer discrimination was established by mixing 10 independently barcoded heterodimers and creating religated barcodes from that mix. 95% of reads pointed to heterodimers, displaying a small underestimation of heterodimers. No hybrid barcodes were seen, showing that no cross-ligation between plasmids or PCR-driven hybrids occur. **F**, A naive barcoding system with no digestion and religation fails because of DNA secondary structures during sequencing.

**Figure S2:**
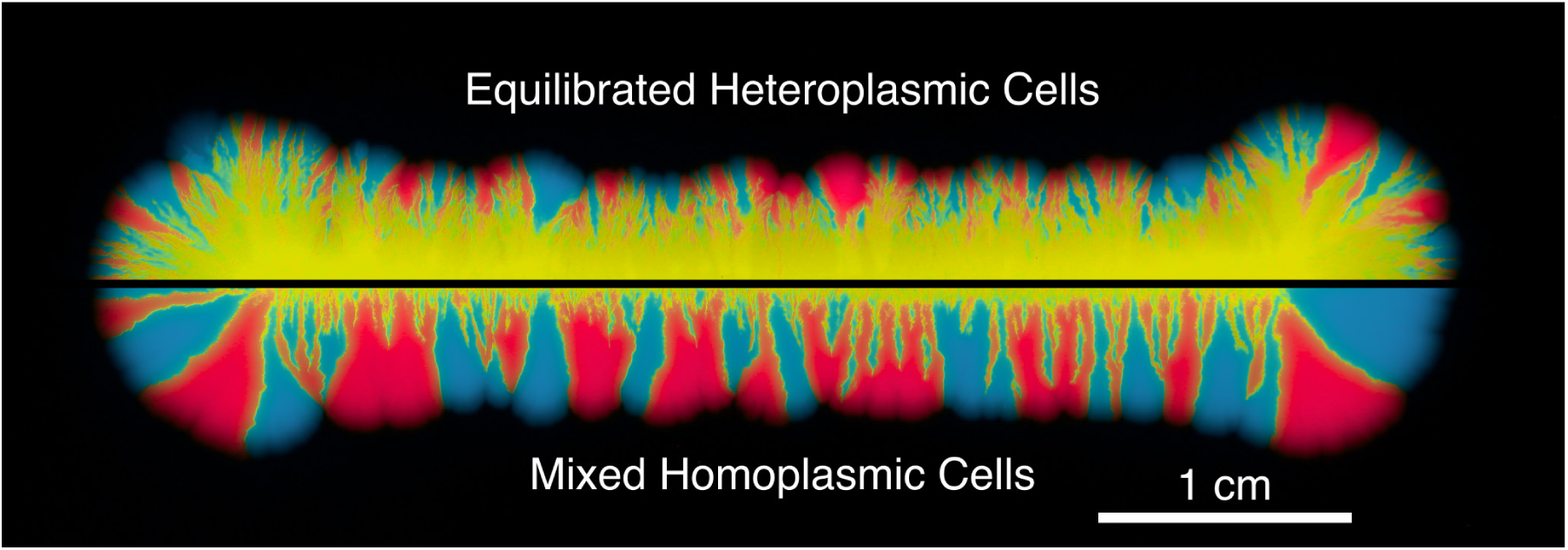
Mixed cells readily segregate as colony expands Comparison between the rapid segregation of a colony expanding from a linear inoculum containing a mix of homoplasmic cells carrying either pScar or pWater (lower half), versus the much slower segregation that occurs when the inoculum cells carried pScar-pWater dimers, which upon FLP induction led to a population of heteroplasmic cells each carrying an equilibrated amount of pScar and pWater (upper half).

**Figure S3:**
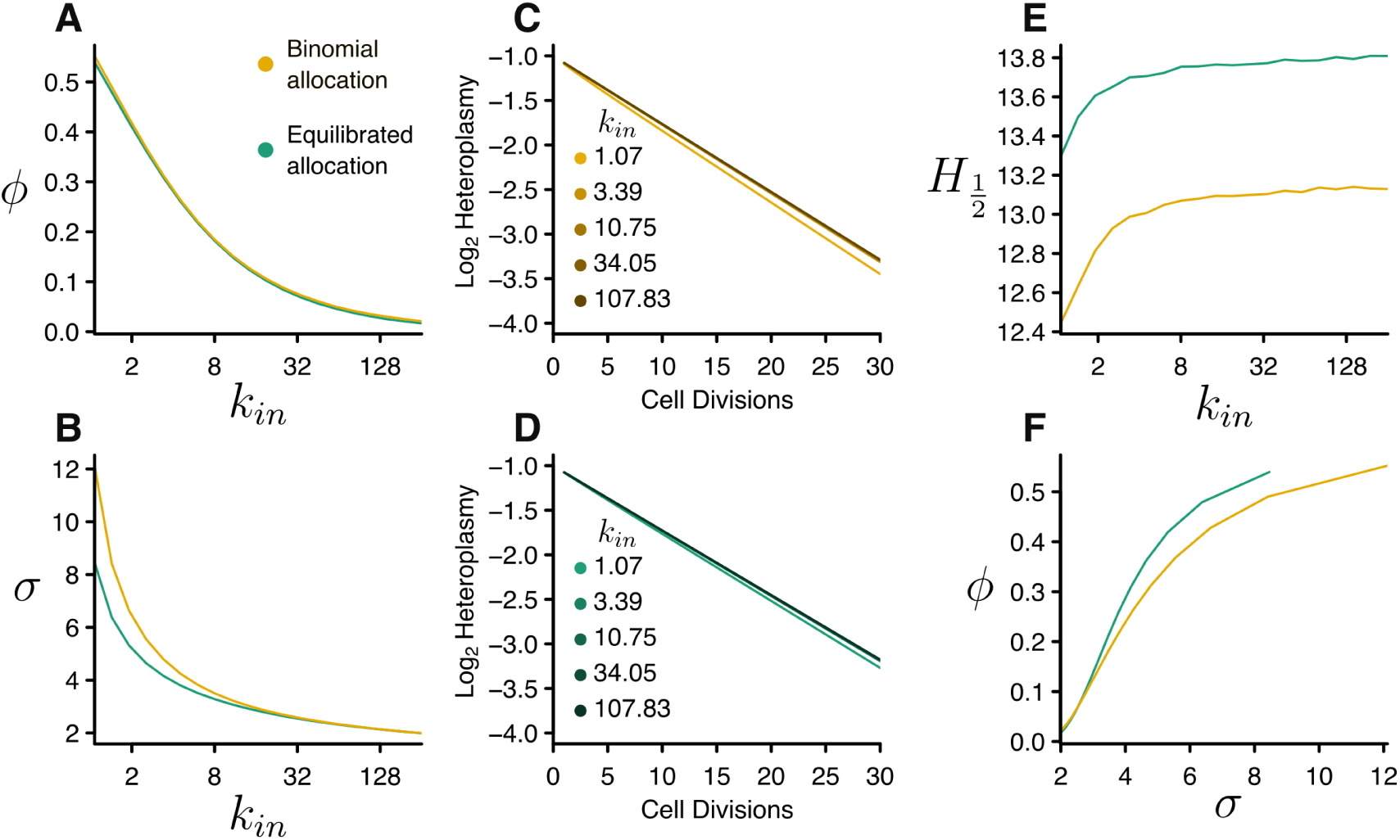
Mechanistic model of plasmid replication imposes strict relationships between parameters Yellow lines show results obtained for model with binomial plasmid partitioning, and green lines for model with equitable partitioning. A,B, Standard deviation *σ* and autocorrelation *ϕ* of stationary distributions are shown for each value of *k_in_*. Stricter replication control decreases variances and makes them less heritable. C,D, Heteroplasmy is shown to decay linearly across conditions. E, modeled heteroplasmy decay rate is much faster than seen empirically.F, Mechanistic model imposes a strict relationship between and *ϕ*.

**Figure S4:**
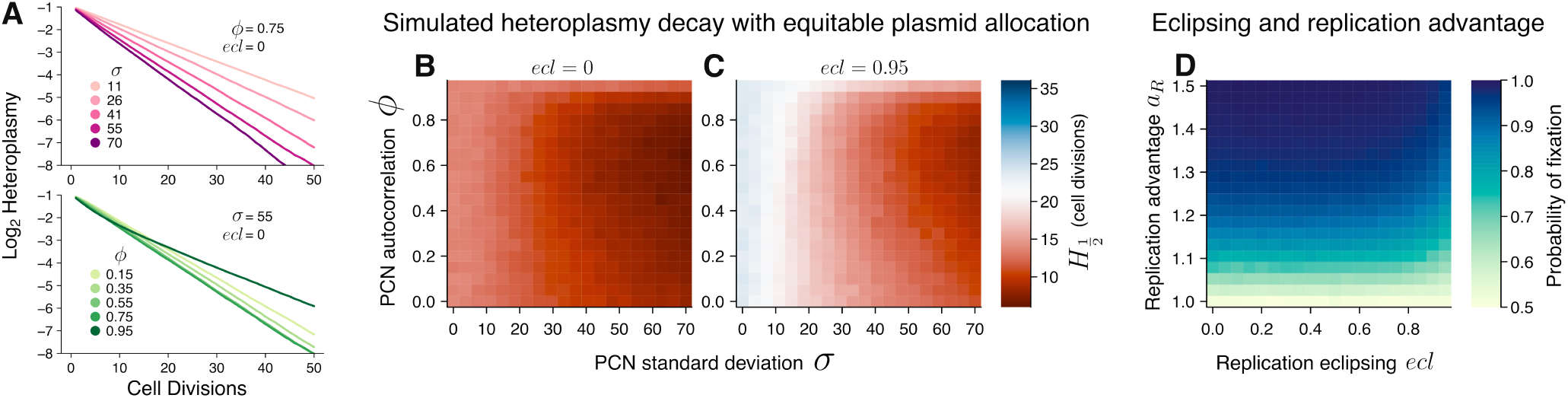
Exponential decay of heteroplasmy **A**, Heteroplasmy decay is shown for a series of values of *σ* and *ϕ*. Higher *σ* accelerates heteroplasmy decay. Conversely, increasing *ϕ* leads to a bi-phasic decay, with an initially fast rate, and a slower asymptotic rate. This bi-phasic behavior occurs because when cells retain similar plasmid copy numbers for many generations there is an initial phase where plasmid fixation quickly occurs across the cell subpopulation that has low copy numbers, while co-occurrence is maintained in the subpopulation with high copy numbers. In a slower time scale, cells with many plasmid copies eventually give rise to cells with few copies, and then fixation occurs in those cells, leading to the second, slower fixation phase. **B,C**, Heteroplasmy decay rates *H* ½ are shown for a modified model in which plasmids are not binomially allocated between daughter cells, but rather each daughter cell receives the same amount of plasmids. This leads to overall slower heteroplasmy decay, but without eclipsing (**B**) still much faster than experimentally observed. **D**, Probability of fixation of a plasmid that has a within-cell competitive advantage is shown as a function of the competitive advantage *a_R_* and the eclipsing factor *ecl*. Higher advantage increases the fixation probability, but stronger eclipsing can diminish the magnitude of the fixation probability increase.

**Figure S5:**
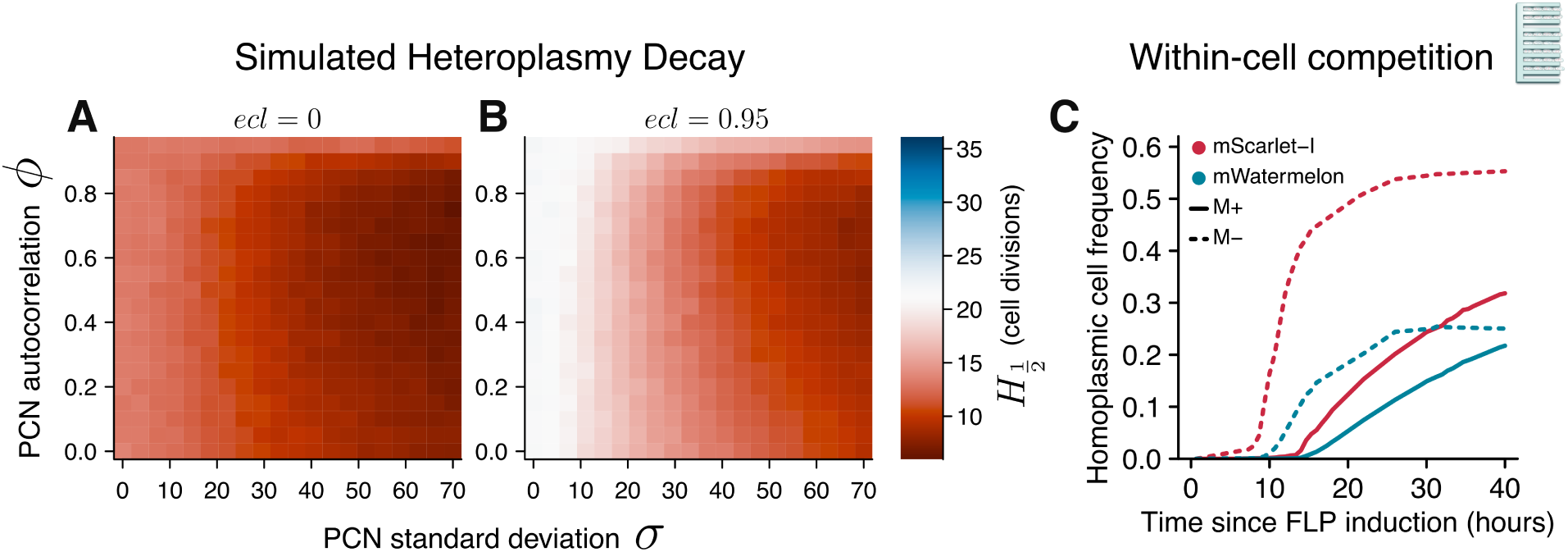
DAM methylation extends within-cell plasmid co-occurrence **A**,**B**, Values of *H* ½ arising from plasmid segregation simulations with (**B**) and without (**A**) eclipsing are shown in red for parameter combinations that lead to faster-than-observed segregation and in blue for slower-than-observed segregation. **C**, Time course of the proportion of cells within a mother machine that fixed one of the competing plasmids. Solid lines represents experiments initiated with cells carrying plasmid dimers with the native methylation sites (pScar-pWater) and dashed lines represent experiments initiated with cells carrying dimers with ablated methylation sites (pScarM^−^-pWaterM^−^).

**Figure S6:**
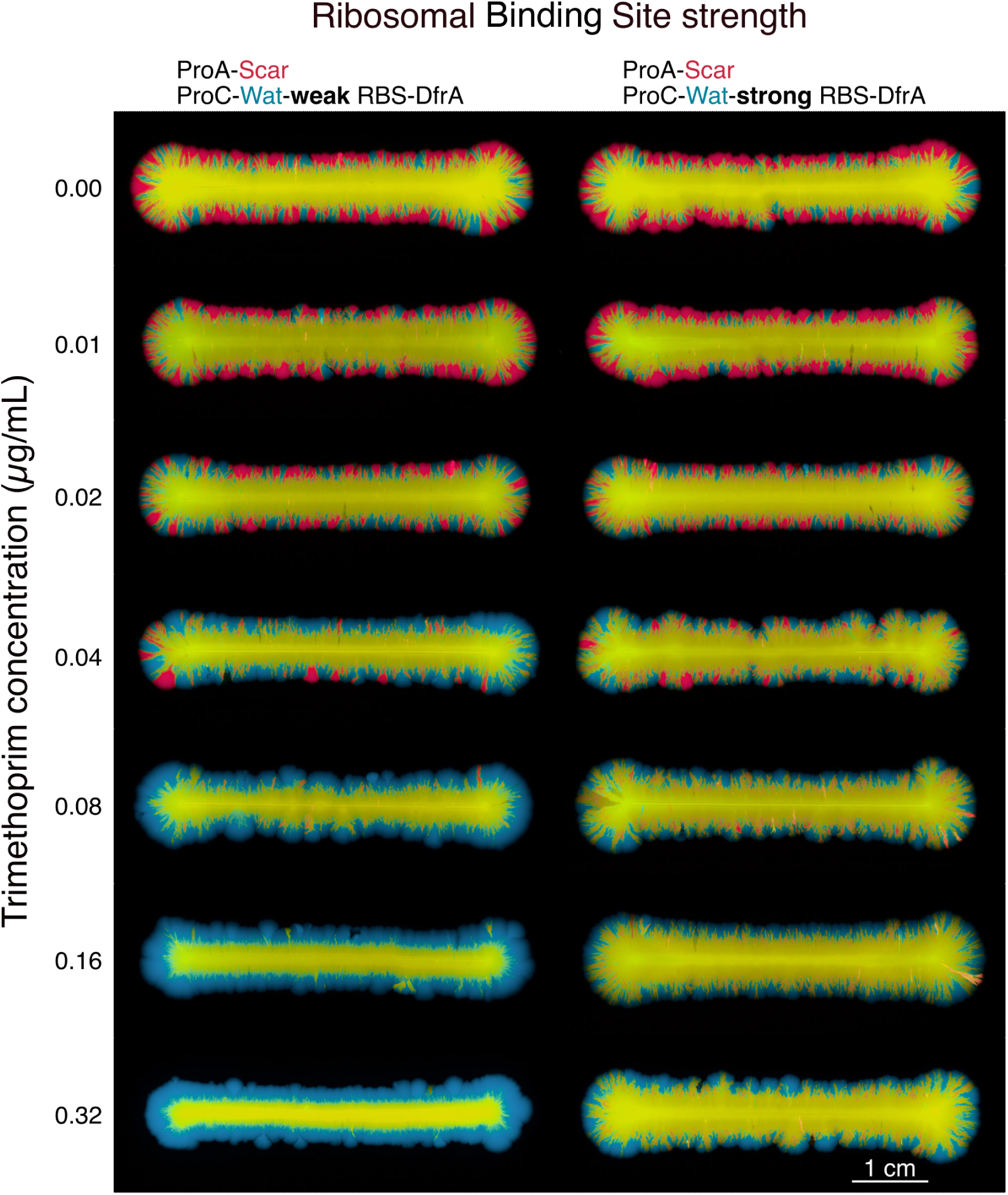
Equilibrated competition experiments Randomly selected samples of competition experiments under various antibiotic concentrations. Cells were inoculated with a razor blade and carried dimers composed of a non-resistance, low-transcription plasmid (shown in red) and a TMP resistance, high-transcription plasmid (shown in blue). Hue denotes the relative normalized brightness of each fluorescent channel.

**Figure S7:**
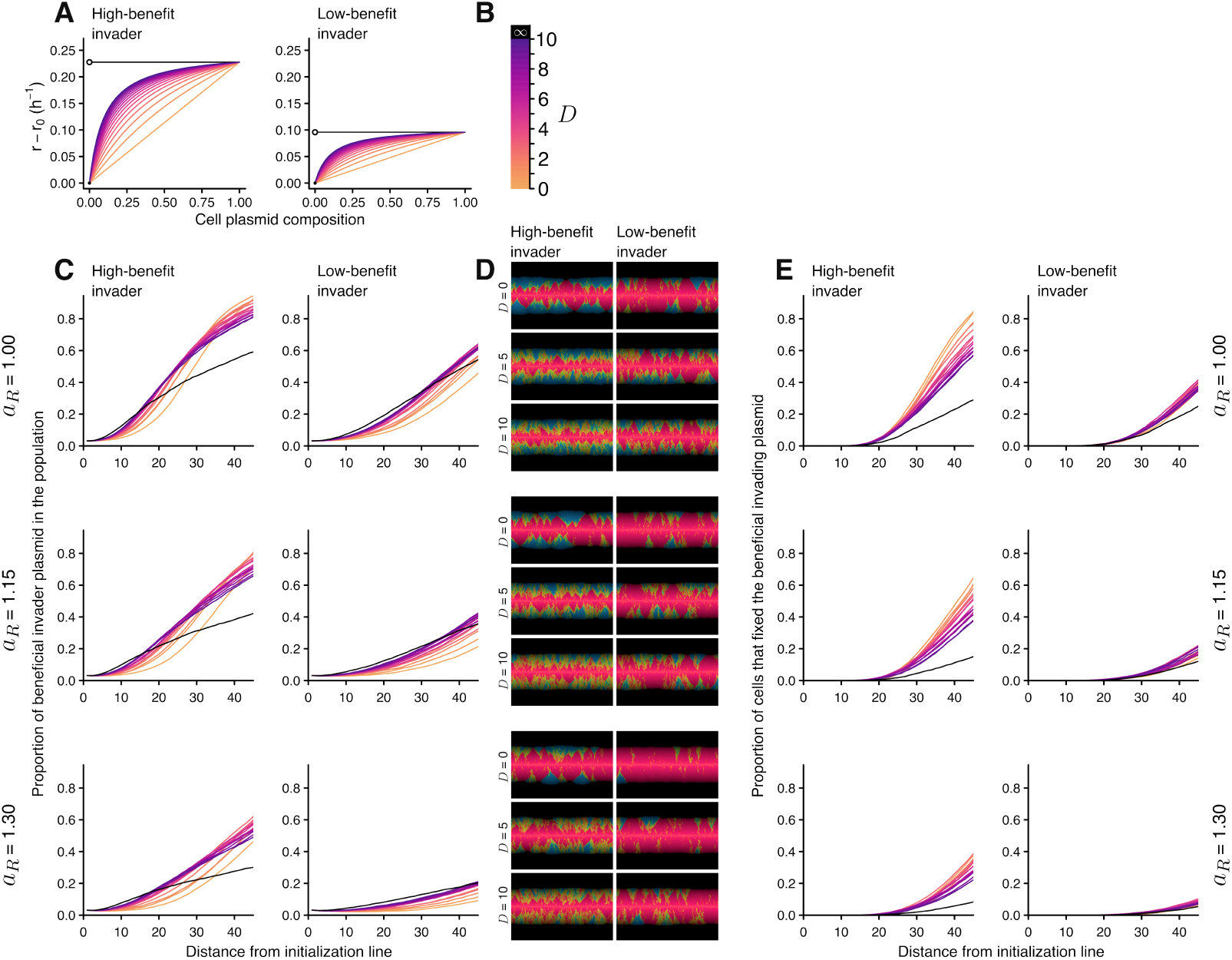
Dominance and within-cell competition at the invasion of a beneficial plasmid **A**, Relative fitness as a function of plasmid composition for cells containing a no-benefit plasmid and either a high benefit plasmid or a low-benefit plasmid. Cell fitness profiles are shown for different potential values of dominance *D*. Black lines show the fitness profile for a completely dominant trait, where any amount of the beneficial plasmid confers the full fitness benefit for the host cell. **B**, Color code for dominance values used throughout the figure. **C**, Average frequency of the invading plasmid as a function of distance from initialization line for highand low-benefit invading plasmids competing with a no-benefit resident plasmid with a within-cell replication advantage given by *a_R_*. Strong dominance always promotes early frequency gains for the beneficial invader. After the initial boost in invader frequency, higher dominance slows subsequent increases of the high-benefit invader, but intermediate dominance still favors the increases of the low-benefit invader. Overall, under stronger within-cell competition (larger *a_R_*), dominance becomes more favorable for the invader. **D**, Simulation samples. The average frequency of invading plasmids can increase by either progressively increasing across cells (yellow areas), or by the appearance of homoplasmic cells carrying only the beneficial plasmid which then expand as a clonal sector (blue areas). In general, higher dominance favors progressive increase and lower dominance favors expansion of homoplasmic sectors. **E**, Spread of homoplasmic (blue) sectors depends on the rate of appearance of homoplasmic cells and the fitness differential between the homoplasmic cells and their surroundings. Increasing dominance can provide an initial frequency boost for the invading plasmid, increasing the stochastic emergence of homoplasmic cells. However, higher dominance also flattens the fitness differential between emergent homoplasmic cells and their surroundings. For high-benefit plasmids, low dominance already allows for frequent emergence of homoplasmic cells, and, therefore, increased dominance only slows the spread of homoplasmic sectors. Conversely, for low-benefit plasmids, intermediate dominance increases the emergence frequency of homoplasmic cells, and overall favors the increase of such sectors.

**Figure S8:**
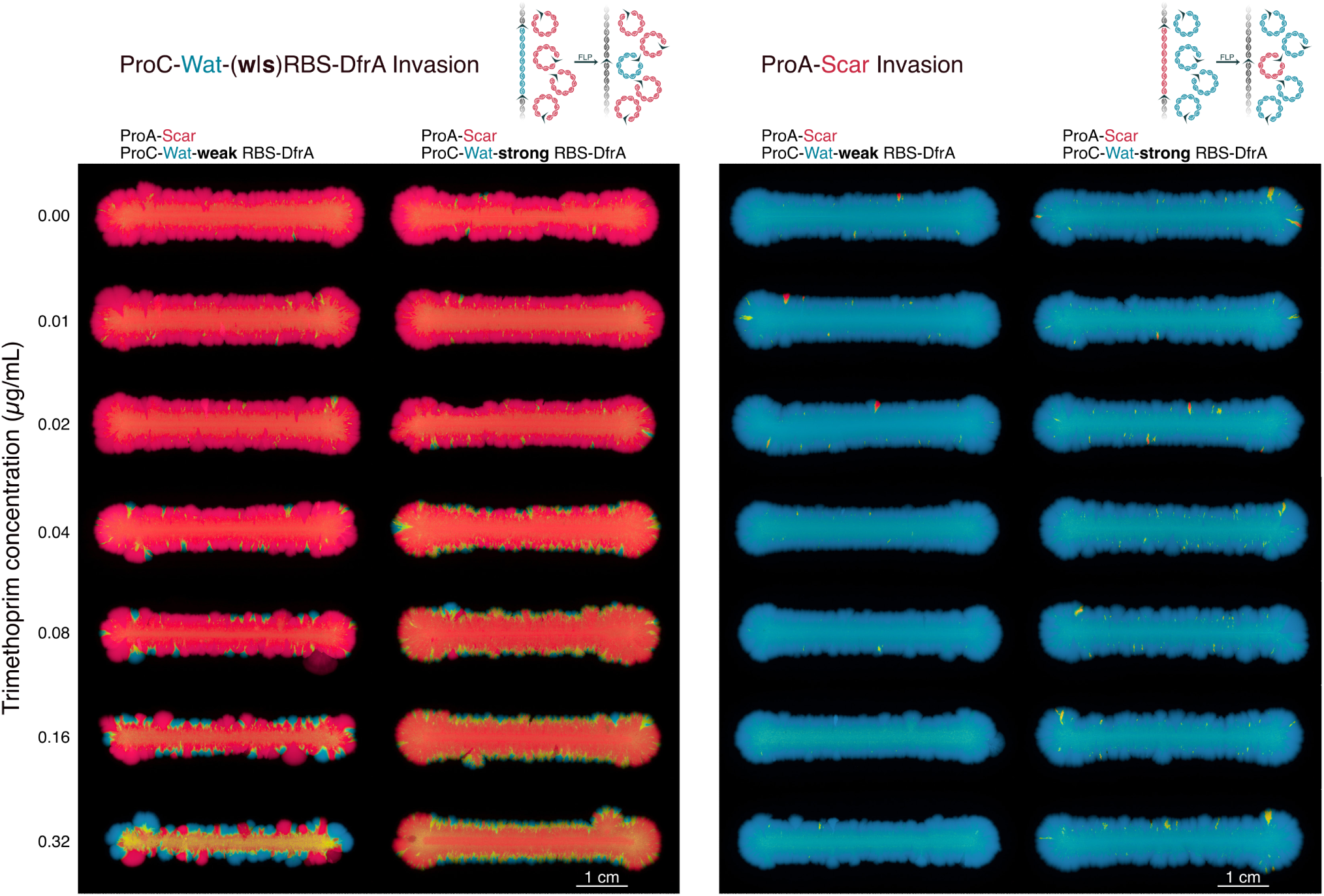
Invasion experiments Randomly selected samples of invasion experiments under various antibiotic concentrations. Cells were inoculated with a razor blade and carried a resident, free plasmid and another plasmid integrated into the chromosome. Upon FLP-induction, the integrated plasmid was excised and started competing with the resident plasmid, approximating a biological invasion. Left panels show experiments for a resident non-resistance, low-transcription plasmid (shown in red) and a chromosomally integrated TMP resistance, high-transcription plasmid (shown in blue). Right panels show experiments for a resident TMP resistance, high-transcription plasmid (shown in blue) and a chromosomally integrated non-resistance, low-transcription plasmid (shown in red).

## Supplemental Movie Captions

**Movie S1:** pScar-pWater competition Time lapse of a razor-blade (3.8cm) inoculation of pScar-pWater dimers 12 hours after inoculation, starting immediately after dimer splitting.

**Movie S2:** Mother machine competition example Mother machine inoculated with a mixture of cells carrying different dimers. First each lane becomes clonal, and then plasmids are split.

**Movie S3:** pScar-pWater competition results in the mother machine

**Movie S4:** Mother machine simulation with mechanistic model Results are similar to mother machine experiments

**Movie S5:** ProC_Water(S)DfrA-ProA_scar competition under 0.04µg/mL TMP Setup is as in Movie S1.

## References

[1] Eörs Szathmáry and John Maynard Smith. The major evolutionary transitions. Nature, 374(6519):227–232, 1995.

[2] Eörs Szathmáry and László Demeter. Group selection of early replicators and the origin of life. Journal of theoretical biology, 128(4):463–486, 1987.

[3] Helen K Matthews, Cosetta Bertoli, and Robertus AM de Bruin. Cell cycle control in cancer. Nature reviews Molecular cell biology, 23(1):74–88, 2022.

[4] Jerónimo Rodríguez-Beltrán, Vidar Sørum, Macarena Toll-Riera, Carmen de la Vega, Rafael Peña-Miller, and Álvaro San Millán. Genetic dominance governs the evolution and spread of mobile genetic elements in bacteria. Proceedings of the National Academy of Sciences, 117(27):15755– 15762, 2020.

[5] Nils F Hülter, Tanita Wein, Johannes Effe, Ana Garoña, and Tal Dagan. Intracellular competitions reveal determinants of plasmid evolutionary success. Frontiers in Microbiology, 11:2062, 2020.

[6] William C Ratcliff, R Ford Denison, Mark Borrello, and Michael Travisano. Experimental evolution of multicellularity. Proceedings of the National Academy of Sciences, 109(5):1595–1600, 2012.

[7] David A Galbraith, Sarah D Kocher, Tom Glenn, Istvan Albert, Greg J Hunt, Joan E Strassmann, David C Queller, and Christina M Grozinger. Testing the kinship theory of intragenomic conflict in honey bees (apis mellifera). Proceedings of the National Academy of Sciences, 113(4):1020–1025, 2016.

[8] Leo W Buss. The evolution of individuality, volume 796. Princeton University Press, 2014.

[9] John H Werren. Selfish genetic elements, genetic conflict, and evolutionary innovation. Proceedings of the National Academy of Sciences, 108(supplement_2):10863–10870, 2011.

[10] Daniel B Cooney. The replicator dynamics for multilevel selection in evolutionary games. Journal of mathematical biology, 79(1):101–154, 2019.

[11] Daniel B Cooney, Fernando W Rossine, Dylan H Morris, and Simon A Levin. A pde model for protocell evolution and the origin of chromosomes via multilevel selection. Bulletin of Mathematical Biology, 84(10):109, 2022.

[12] Frances Medaney, Richard J Ellis, and Ben Raymond. Ecological and genetic determinants of plasmid distribution in e scherichia coli. Environmental Microbiology, 18(11):4230–4239, 2016.

[13] Emily C Fogarty, Matthew S Schechter, Karen Lolans, Madeline L Sheahan, Iva Veseli, Ryan M Moore, Evan Kiefl, Thomas Moody, Phoebe A Rice, K Yu Michael, et al. A cryptic plasmid is among the most numerous genetic elements in the human gut. Cell, 187(5):1206–1222, 2024.

[14] Michael K Yu, Emily C Fogarty, and A Murat Eren. Diverse plasmid systems and their ecology across human gut metagenomes revealed by plasx and mobmess. Nature Microbiology, 9(3):830– 847, 2024.

[15] Manuel Ares-Arroyo, Eduardo PC Rocha, and Bruno Gonzalez-Zorn. Evolution of cole1-like plasmids across *γ*-proteobacteria: From bacteriocin production to antimicrobial resistance. PLoS Genetics, 17(11):e1009919, 2021.

[16] Barbara E Funnell and Gregory J Phillips. Plasmid biology, volume 672. ASM press Washington, DC, 2004.

[17] Patricia L Shipley, Alfred D Allen, and Thomas N Swanson. Cointegrate formation between plasmids carrying virulence factor and antibiotic resistance genes in e. coli. FEMS microbiology letters, 20(3):365–368, 1983.

[18] Barbara Froehlich, Julian Parkhill, Mandy Sanders, Michael A Quail, and June R Scott. The pcoo plasmid of enterotoxigenic escherichia coli is a mosaic cointegrate. Journal of bacteriology, 187(18):6509–6516, 2005.

[19] Aurora García-Fernández, Daniela Fortini, Kees Veldman, Dik Mevius, and Alessandra Carattoli. Characterization of plasmids harbouring qnrs1, qnrb2 and qnrb19 genes in salmonella. Journal of Antimicrobial Chemotherapy, 63(2):274–281, 2009.

[20] M Rosario Rodicio, Ana Herrero, Irene Rodríguez, Patricia García, Ignacio Montero, Janine Beutlich, Rosaura Rodicio, Beatriz Guerra, and M Carmen Mendoza. Acquisition of antimicrobial resistance determinants by virulence plasmids specific for nontyphoid serovars of salmonella enterica. Reviews and Research in Medical Microbiology, 22(3):55–65, 2011.

[21] Haiyang Liu, Yuexing Tu, Jintao He, Qingye Xu, Xiaofan Zhang, Xinli Mu, Minhua Chen, Hua Zhou, and Xi Li. Emergence and plasmid cointegration-based evolution of ndm-1-producing st107 citrobacter freundii high-risk resistant clone in china. International Journal of Antimicrobial Agents, 63(2):107069, 2024.

[22] Sean Conlan, Pamela J Thomas, Clayton Deming, Morgan Park, Anna F Lau, John P Dekker, Evan S Snitkin, Tyson A Clark, Khai Luong, Yi Song, et al. Single-molecule sequencing to track plasmid diversity of hospital-associated carbapenemase-producing enterobacteriaceae. Science translational medicine, 6(254):254ra126–254ra126, 2014.

[23] Santiago Redondo-Salvo, Raúl Fernández-López, Raúl Ruiz, Luis Vielva, María de Toro, Eduardo PC Rocha, M Pilar Garcillán-Barcia, and Fernando de la Cruz. Pathways for horizontal gene transfer in bacteria revealed by a global map of their plasmids. Nature communications, 11(1):3602, 2020.

[24] Holger Heuer and Kornelia Smalla. Plasmids foster diversification and adaptation of bacterial populations in soil. FEMS microbiology reviews, 36(6):1083–1104, 2012.

[25] Alvaro San Millan, Jose Antonio Escudero, Danna R Gifford, Didier Mazel, and R Craig MacLean. Multicopy plasmids potentiate the evolution of antibiotic resistance in bacteria. Nature ecology & evolution, 1(1):0010, 2016.

[26] Olivia Kosterlitz, Nathan Grassi, Bailey Werner, Ryan Seamus McGee, Eva M Top, and Benjamin Kerr. Evolutionary “crowdsourcing”: alignment of fitness landscapes allows for cross-species adaptation of a horizontally transferred gene. Molecular Biology and Evolution, 40(11):msad237, 2023.

[27] Richard P Novick. Plasmid incompatibility. Microbiological reviews, 51(4):381–395, 1987.

[28] Shalni Kumar, Andrew Lezia, and Jeff Hasty. Engineering plasmid copy number heterogeneity for dynamic microbial adaptation. Nature Microbiology, pages 1–12, 2024.

[29] Stéphanie Bedhomme, D Perez Pantoja, and Ignacio G Bravo. Plasmid and clonal interference during post horizontal gene transfer evolution. Molecular ecology, 26(7):1832–1847, 2017.

[30] Ana Garoña, Mario Santer, Nils F Hülter, Hildegard Uecker, and Tal Dagan. Segregational drift hinders the evolution of antibiotic resistance on polyploid replicons. PLoS Genetics, 19(8):e1010829, 2023.

[31] Judith Ilhan, Anne Kupczok, Christian Woehle, Tanita Wein, Nils F Hülter, Philip Rosenstiel, Giddy Landan, Itzhak Mizrahi, and Tal Dagan. Segregational drift and the interplay between plasmid copy number and evolvability. Molecular biology and evolution, 36(3):472–486, 2019.

[32] Michael R Snaith, Nigel J Kilby, and James AH Murray. An escherichia coli system for assay of flp site-specific recombination on substrate plasmids. Gene, 180(1-2):225–227, 1996.

[33] DK Summers and DJ Sherratt. Resolution of cole1 dimers requires a dna sequence implicated in the three-dimensional organization of the cer site. The EMBO Journal, 7(3):851–858, 1988.

[34] Diego Libkind, Chris Todd Hittinger, Elisabete Valério, Carla Gonçalves, Jim Dover, Mark Johnston, Paula Gonçalves, and José Paulo Sampaio. Microbe domestication and the identification of the wild genetic stock of lager-brewing yeast. Proceedings of the National Academy of Sciences, 108(35):14539–14544, 2011.

[35] Daphne S Bindels, Lindsay Haarbosch, Laura Van Weeren, Marten Postma, Katrin E Wiese, Marieke Mastop, Sylvain Aumonier, Guillaume Gotthard, Antoine Royant, Mark A Hink, et al. mscarlet: a bright monomeric red fluorescent protein for cellular imaging. Nature methods, 14(1):53–56, 2017.

[36] Khyati Gohil, Sheng-Yi Wu, Kei Takahashi-Yamashiro, Yi Shen, and Robert E Campbell. Biosensor optimization using a forster resonance energy transfer pair based on mscarlet red fluorescent protein and an mscarlet-derived green fluorescent protein. ACS sensors, 8(2):587–597, 2023.

[37] Oskar Hallatschek, Pascal Hersen, Sharad Ramanathan, and David R Nelson. Genetic drift at expanding frontiers promotes gene segregation. Proceedings of the National Academy of Sciences, 104(50):19926–19930, 2007.

[38] Ping Wang, Lydia Robert, James Pelletier, Wei Lien Dang, Francois Taddei, Andrew Wright, and Suckjoon Jun. Robust growth of escherichia coli. Current biology, 20(12):1099–1103, 2010.

[39] Nathalie Q Balaban, Jack Merrin, Remy Chait, Lukasz Kowalik, and Stanislas Leibler. Bacterial persistence as a phenotypic switch. Science, 305(5690):1622–1625, 2004.

[40] Laurent Potvin-Trottier, Scott Luro, and Johan Paulsson. Microfluidics and single-cell microscopy to study stochastic processes in bacteria. Current opinion in microbiology, 43:186–192, 2018.

[41] Jeffrey E Barrick, Dong Su Yu, Sung Ho Yoon, Haeyoung Jeong, Tae Kwang Oh, Dominique Schneider, Richard E Lenski, and Jihyun F Kim. Genome evolution and adaptation in a long-term experiment with escherichia coli. Nature, 461(7268):1243–1247, 2009.

[42] Johan Paulsson and Måns Ehrenberg. Noise in a minimal regulatory network: plasmid copy number control. Quarterly reviews of biophysics, 34(1):1–59, 2001.

[43] Pradeep K Patnaik, S Merlin, and BARRY Polisky. Effect of altering gatc sequences in the plasmid cole1 primer promoter. Journal of bacteriology, 172(4):1762–1768, 1990.

[44] Agnes Bergerat, Anastasios Kriebardis, and Wilhelm Guschlbauer. Preferential site-specific hemimethylation of gatc sites in pbr322 dna by dam methyltransferase from escherichia coli. Journal of Biological Chemistry, 264(7):4064–4070, 1989.

[45] Daghfous Douraid and Landoulsi Ahmed. Seqa, the escherichia coli origin sequestration protein, can regulate the replication of the pbr322 plasmid. Plasmid, 65(1):15–19, 2011.

[46] Motoo Kimura and Tomoko Ohta. The average number of generations until fixation of a mutant gene in a finite population. Genetics, 61(3):763, 1969.

[47] Houra Merrikh, Yan Zhang, Alan D Grossman, and Jue D Wang. Replication–transcription conflicts in bacteria. Nature Reviews Microbiology, 10(7):449–458, 2012.

[48] Leroy F Liu and James C Wang. Supercoiling of the dna template during transcription. Proceedings of the National Academy of Sciences, 84(20):7024–7027, 1987.

[49] Joseph H Davis, Adam J Rubin, and Robert T Sauer. Design, construction and characterization of a set of insulated bacterial promoters. Nucleic acids research, 39(3):1131–1141, 2011.

[50] Howard M Salis, Ethan A Mirsky, and Christopher A Voigt. Automated design of synthetic ribosome binding sites to control protein expression. Nature biotechnology, 27(10):946–950, 2009.

[51] Richard Durrett and Simon Levin. The importance of being discrete (and spatial). Theoretical population biology, 46(3):363–394, 1994.

[52] Michael Baym, Tami D Lieberman, Eric D Kelsic, Remy Chait, Rotem Gross, Idan Yelin, and Roy Kishony. Spatiotemporal microbial evolution on antibiotic landscapes. Science, 353(6304):1147– 1151, 2016.

[53] James F Crow and Motoo Kimura. Evolution in sexual and asexual populations. The American Naturalist, 99(909):439–450, 1965.

[54] Eörs Szathmáry. Toward major evolutionary transitions theory 2.0. Proceedings of the National Academy of Sciences, 112(33):10104–10111, 2015.

[55] Ellie Harrison and Michael A Brockhurst. Plasmid-mediated horizontal gene transfer is a coevolutionary process. Trends in microbiology, 20(6):262–267, 2012.

[56] Rajiv I Modi and Julian Adams. Coevolution in bacterial-plasmid populations. Evolution, 45(3):656– 667, 1991.

[57] AJ Spiers and PL Bergquist. Expression and regulation of the repa protein of the repfib replicon from plasmid p307. Journal of bacteriology, 174(23):7533–7541, 1992.

[58] Nalini Raghunathan, Sayantan Goswami, Jakku K Leela, Apuratha Pandiyan, and Jayaraman Gowrishankar. A new role for escherichia coli dam dna methylase in prevention of aberrant chromosomal replication. Nucleic acids research, 47(11):5698–5711, 2019.

[59] AL Abeles and SJ Austin. P1 plasmid replication requires methylated dna. The EMBO Journal, 6(10):3185–3189, 1987.

[60] Sandra Martin, Florian Fournes, Giovanna Ambrosini, Christian Iseli, Karolina Bojkowska, Julien Marquis, Nicolas Guex, and Justine Collier. Dna methylation by ccrm contributes to genome maintenance in the agrobacterium tumefaciens plant pathogen. Nucleic Acids Research, 52(19):11519– 11535, 2024.

[61] David Forrest, Emily A Warman, Amanda M Erkelens, Remus T Dame, and David C Grainger. Xenogeneic silencing strategies in bacteria are dictated by rna polymerase promiscuity. Nature communications, 13(1):1149, 2022.

[62] William Wiley Navarre. H-ns as a defence system. Bacterial chromatin, pages 251–322, 2010.

[63] Jerónimo Rodríguez-Beltrán, Ricardo León-Sampedro, Paula Ramiro-Martínez, Carmen de la Vega, Fernando Baquero, Bruce R Levin, and Álvaro San Millán. Translational demand is not a major source of plasmid-associated fitness costs. Philosophical Transactions of the Royal Society B, 377(1842):20200463, 2022.

[64] Alvaro San Millan, Macarena Toll-Riera, Qin Qi, Alex Betts, Richard J Hopkinson, James McCullagh, and R Craig MacLean. Integrative analysis of fitness and metabolic effects of plasmids in pseudomonas aeruginosa pao1. The ISME journal, 12(12):3014–3024, 2018.

[65] Stephen Fitzgerald, Stefani C Kary, Ebtihal Y Alshabib, Keith D MacKenzie, Daniel M Stoebel, TzuChiao Chao, and Andrew DS Cameron. Redefining the h-ns protein family: a diversity of specialized core and accessory forms exhibit hierarchical transcriptional network integration. Nucleic Acids Research, 48(18):10184–10198, 2020.

[66] Eleanor A Rand, Sian V Owen, Natalia Quinones-Olvera, Kesther Jean, Carmen Hernandez-Perez, and Michael Baym. Phage disco: targeted discovery of bacteriophages by co-culture. bioRxiv, pages 2024–11, 2024.

[67] Daniel Eaton, Carlos Sanchez, Luis Alberto Gutierrez-Lopez, Jacob Shenker, Youlian Goulev, Quincey Justman, Brianna Watson, Veronique Henriot, Ethan Garner, Jeff Moffit, and Johan Paulsson. Multigenerational optical pooled screen reveals growth control by elongation sensing. in prep, 2025.

[68] Anders Løbner-Olesen. Distribution of minichromosomes in individual escherichia coli cells: implications for replication control. The EMBO journal, 1999.

[69] Bin Shao, Jayan Rammohan, Daniel A Anderson, Nina Alperovich, David Ross, and Christopher A Voigt. Single-cell measurement of plasmid copy number and promoter activity. Nature communications, 12(1):1475, 2021.

[70] Shay Tal and Johan Paulsson. Evaluating quantitative methods for measuring plasmid copy numbers in single cells. Plasmid, 67(2):167–173, 2012.

[71] Matteo Osella, Eileen Nugent, and Marco Cosentino Lagomarsino. Concerted control of escherichia coli cell division. Proceedings of the National Academy of Sciences, 111(9):3431–3435, 2014.

