## Supplementary material for "Intracellular competition shapes plasmid population dynamics": DNA Sequences of synthetic constructs

Auxiliary plasmid:

GAGTTATACACAGGGCTGGGATCTATTCTTTTTATCTTTTTTTATTCTTTCTTTATTCTATAAATTATA
ACCACTTGAATATAAACAAAAAAAACACACAAAGGTCTAGCGGAATTTACAGAGGGTCTAGCAGAATTT
ACAAGTTTTCCAGCAAAGGTCTAGCAGAATTTACAGATACCCACAACTCAAAGGAAAAGGACTAGTAAT
TATCATTGACTAGCCCATCTCAATTGGTATAGTGATTAAAATCACCTAGACCAATTGAGATGTATGTCT
GAATTAGTTGTTTTCAAAGCAAATGAACTAGCGATTAGTCGCTATGACTTAACGGAGCATGAAACCAAG
CTAATTTTATGCTGTGTGGCACTACTCAACCCCACGATTGAAAACCCTACAAGGAAAGAACGGACGGTA
TCGTTCACTTATAACCAATACGCTCAGATGATGAACATCAGTAGGGAAAATGCTTATGGTGTATTAGCT
AAAGCAACCAGAGAGCTGATGACGAGAACTGTGGAAATCAGGAATCCTTTGGTTAAAGGCTTTGAGATT
TTCCAGTGGACAAACTATGCCAAGTTCTCAAGCGAAAAATTAGAATTAGTTTTTAGTGAAGAGATATTG
CCTTATCTTTTCCAGTTAAAAAAATTCATAAAATATAATCTGGAACATGTTAAGTCTTTTGAAAACAAA
TACTCTATGAGGATTTATGAGTGGTTATTAAAAGAACTAACACAAAAGAAAACTCACAAGGCAAATATA
GAGATTAGCCTTGATGAATTTAAGTTCATGTTAATGCTTGAAAATAACTACCATGAGTTTAAAAGGCTT
AACCAATGGGTTTTGAAACCAATAAGTAAAGATTTAAACACTTACAGCAATATGAAATTGGTGGTTGAT
AAGCGAGGCCGCCCGACTGATACGTTGATTTTCCAAGTTGAACTAGATAGACAAATGGATCTCGTAACC
GAACTTGAGAACAACCAGATAAAAATGAATGGTGACAAAATACCAACAACCATTACATCAGATTCCTAC
CTACATAACGGACTAAGAAAAACACTACACGATGCTTTAACTGCAAAAATTCAGCTCACCAGTTTTGAG
GCAAAATTTTTGAGTGACATGCAAAGTAAGTATGATCTCAATGGTTCGTTCTCATGGCTCACGCAAAAA
CAACGAACCACACTAGAGAACATACTGGCTAAATACGGAAGGATCTGAGGTTCTTATGGCTCTTGTATC
TATCAGTGAAGCATCAAGACTAACAAACAAAAGTAGAACAACTGTTCACCGTTACATATCAAAGGGAAA
ACTGTCCATATGCACAGATGAAAACGGTGTAAAAAAGATAGATACATCAGAGCTTTTACGAGTTTTTGG
TGCATTCAAAGCTGTTCACCATGAACAGATCGACAATGTAACAGATGAACAGCATGTAACACCTAATAG
AACAGGTGAAACCAGTAAAACAAAGCAACTAGAACATGAAATTGAACACCTGAGACAACTTGTTACAGC
TCAACAGTCACACATAGACAGCCTGAAACAGGCGATGCTGCTTATCGAATCAAAGCTGCCGACAACACG
GGAGCCAGTGACGCCTCCCGTGGGGAAAAAATCATGGCAATTCTGGAAGAAATAGCGCTTTCAGCCGGC
AAACCGGCTGAAGCCGGATCTGCGATTCTGATAACAAACTAGCAACACCAGAACAGCCCGTTTGCGGGC
AGCAAAACCCGTACCCTAGGTCTAGGGCGGCGGATTTGTCCTACTCAGGAGAGCGTTCACCGACAAACA
ACAGATAAAACGAAAGGCCCAGTCTTTCGACTGAGCCTTTCGTTTTATTTGATGCCTCTAGAGCTTGCA
TGCCTGCAGGTTAAATGCGGCGGCTAATATAGCTGCTCAGATAATCCAGCACTTCTTGCGCAATAATGC
CGTTCCACGCCGCATAGCGGGTGCTGCCGCCGGTGCTGCCTTTCAGCTGTTCAATATGGCGCCATTCTT
CAATCGGGTTGGTTTCATCTTTCCACGGAATCATTTCTTTGCTAATCTGATCATAGCCATAATAGCCGC
TCACCAGCGCAAAATAATGATCCGGAATCGCGGTCACTTGATGGGTATAGGTGGTGCGCGCCACCACGC
TCGCGCGTTTATCGCTCCAGTTGCCCACCACGTTGGTCAGTTCGGTCAGGCTTTTCATGCTCAGAAAGC
TCGTCATCAGATGGCGGCCAATATGGCTTTTCGGGCCGTTTTTAATCGCCAGAATGCTATACGGCGCGT
TGCTTTTCAGCGCTTTGTTATAGCTGCGCACCAGGTTATCTTTCAGCAGCTGATACTGCTGTTTGTTGC
CGCTGCTGCTGCTCGTTTTGTTAATGCGTTTCGGCACCGGTTCGCTATAGCGCAGAAATTCATCCAGAT
ACACCAGGCTATCCAGGCGGCCTTTCGCGCTAAAAAAATAAATATGGCGGCTCACGCCGGTTTTGGTTT
CGGTCACCAGGCACTGAATAATCACGCCCAGATATTCGTTCTGAATCAGTTTAAAGCTCTGCGGATCCA
CGTTTTTAATATCGCTAAAGCGCGCGCAGTTCACAAAGGTCGCCAGAAACAGAAACTGATACTGCGCTT
TGGTTTTGGTATAGCGGCTCGTATATTCAAAGCTATCCAGAATTTTTTCCGCAATGTTCCACACGCTTT
CATCTTCGTTCAGCAGCGCTTTCAGCATTTTTTTGCTATGGCTGTTGCCTTTTTCCACTTCTTCCGGGC
TTTCAAACTGCAGCTGCAGGTTGCTCACAATATCGGTCACATCGCTCTGTTCTTTCTGACCATAATACG
GAATAATGGTAAATTCCCAGCCCGGGATCAGCTTCTGCAGGCTCGCTTGCAGAATCGCCGCTTTCTGCG
TTTTATATTTAAACTGCAGGGTTTTTTTCACCACATCATACTGCAGGCTTTTGCTAATAATGGTGTTAT

AGCTCAGAAAGGTCGCGCGTTTAATCGCCGCGCCGTTATGGGTAATCATCCAGCACAGATAGGTCAGTT
CCGCCGCGCAGCTCGCCAGTTTTTCGCCGCTCGGTTCGCCAAAGCGCGCAATAAACTGACTCACCAGCA
CTTTCGGCGGGGTTTTATACAGAATATCAAATTTGCTCATGGTGAATTCCTCCTGCTAGCCCAAAAAAA
CGGGTATGGAGAAACAGTAGAGAGTTGCGATAAAAAGCGTCAGGTAGGATCCGCTAATCTTATGGATAA
AAATGCTATGGCATAGCAAAGTGTGACGCCGTGCAAATAATCAATGTGGACTTTTCTGCCGTGATTATA
GACACTTTTGTTACGCGTTTTTGTCATGGCTTTGGTCCCGCTTTGTTACAGAATGCTTTTAATAAGCGG
GGTTACCGGTTTGGTTAGCGAGAAGAGCCAGTAAAAGACGCAGTGACGGCAATGTCTGATGCAATATGG
ACAATTGGTTTCTTCTCTGAATGGCGGGAGTATGAAAAGTATGGCTGAAGCGCAAAATGATCCCCTGCT
GCCGGGATACTCGTTTAATGCCCATCTGGTGGCGGGTTTAACGCCGATTGAGGCCAACGGTTATCTCGA
TTTTTTTATCGACCGACCGCTGGGAATGAAAGGTTATATTCTCAATCTCACCATTCGCGGTCAGGGGGT
GGTGAAAAATCAGGGACGAGAATTTGTTTGCCGACCGGGTGATATTTTGCTGTTCCCGCCAGGAGAGAT
TCATCACTACGGTCGTCATCCGGAGGCTCGCGAATGGTATCACCAGTGGGTTTACTTTCGTCCGCGCGC
CTACTGGCATGAATGGCTTAACTGGCCGTCAATATTTGCCAATACGGGGTTCTTTCGCCCGGATGAAGC
GCACCAGCCGCATTTCAGCGACCTGTTTGGGCAAATCATTAACGCCGGGCAAGGGGAAGGGCGCTATTC
GGAGCTGCTGGCGATAAATCTGCTTGAGCAATTGTTACTGCGGCGCATGGAAGCGATTAACGAGTCGCT
CCATCCACCGATGGATAATCGGGTACGCGAGGCTTGTCAGTACATCAGCGATCACCTGGCAGACAGCAA
TTTTGATATCGCCAGCGTCGCACAGCATGTTTGCTTGTCGCCGTCGCGTCTGTCACATCTTTTCCGCCA
GCAGTTAGGGATTAGCGTCTTAAGCTGGCGCGAGGACCAACGTATCAGCCAGGCGAAGCTGCTTTTGAG
CACCACCCGGATGCCTATCGCCACCGTCGGTCGCAATGTTGGTTTTGACGATCAACTCTATTTCTCGCG
GGTATTTAAAAAATGCACCGGGGCCAGCCCGAGCGAGTTCCGTGCCGGTTGTGAAGAAAAAGTGAATGA
TGTAGCCGTCAAGTTGTCATAATTGGTAACGAATCAGACAATTGACGGCTTGACGGAGTAGCATAGGGT
TTGCAGAATCCCTGCTTCGTCCATTTGACAGGCACATTATGCATGCCCGTAAAGTTATCCAGCAACCAC
TCATAGACCTAGGGCAGCAGATAGGGACGACGTGGTGTTAGCTGTGGTGAAGACGAAAGGCCTCGTGA
TACGCCTATTTTTATAGGTTAATGTCATGATAATAATGGTTTCTTAGACGTCGGAATTGCCAGCTGGTT
AATTAAGGTTTCTTAGACGTCAGGTGGCACTTTGACTGGAGTTCAGACGTGTGCTCTTCCGATCTGTGG
GTACCGCGCCCTCTGGTAAGTGATCGGCACGTAAGAGGTTCCAACTTTCACCATAATGAAATAAGATCA
CTACCGGGCGTATTTTTTGAGTTGTCGAGATTTTCAGGAGCTAAGGAAGCTAAAATGGAGAAAAAAATC
ACTGGATATACCACCGTTGATATATCCCAATGGCATCGTAAAGAACATTTTGAGGCATTTCAGTCAGTT
GCTCAATGTACCTATAACCAGACCGTTCAGCTGGATATTACGGCCTTTTTAAAGACCGTAAAGAAAAAT
AAGCACAAGTTTTATCCGGCCTTTATTCACATTCTTGCCCGCCTGATGAATGCTCATCCGGAATTACGT
ATGGCAATGAAAGACGGTGAGCTGGTGATATGGGATAGTGTTCACCCTTGTTACACCGTTTTCCATGAG
CAAACTGAAACGTTTTCATCGCTCTGGAGTGAATACCACGACGATTTCCGGCAGTTTCTACACATATAT
TCGCAAGATGTGGCGTGTTACGGTGAAAACCTGGCCTATTTCCCTAAAGGGTTTATTGAGAATATGTTT
TTCGTCTCAGCCAATCCCTGGGTGAGTTTCACCAGTTTTGATTTAAACGTGGCCAATATGGACAACTTC
TTCGCCCCCGTTTTCACCATGGGCAAATATTATACGCAAGGCGACAAGGTGCTGATGCCGCTGGCGATT
CAGGTTCATCATGCCGTTTGTGATGGCTTCCATGTCGGCAGAATGCTTAATGAATTACAACAGTACTGC
GATGAGTGGCAGGGCGGGGCGTAATAAGGACTCTGGGATCTCTGCAGTCGCGATGATTAATTAATTCAG
AACGCTCGGTTGCCGCCGGGCGTTTTTTATGCATGAGAATCCTGGCGGGTCTGTCGACGCACTAGGGAC
AGTAAGACGGGTAAGCCTGTTGATGATACCGCTGCCTTACTGGGTGCATTAGCCAGTCTGAATGACCTG
TCACGGGATAATCCGAAGTGGTCAGACTGGAAAATCAGAGGGCAGGAACTGCTGAACAGCAAAAAGTCA
GATAGCACCACATAGCAGACCCGCCATAAAACGCCCTGAGAAGCCCGTGACGGGCTTTTCTTGTATTAT
GGGTAGTTTCCTTGCATGAATCCATAAAAGGCGCCTGTAGTGCCATTTACCCCCATTCACTGCCAGAGC
CGTGAGCGCAGCGAACTGAATGTCACGAAAAAGACAGCGACTCAGGTGCCTGATGGTCGGAGACAAAAG
GAATATTCAGCGATTTGCCCGAGCTTGCGAGGGTGCTACTTAAGCCTTTAGGGTTTTAAGGTCTGTTTT
GTAGAGGAGCAAACAGCGTTTGCGACATCCTTTTGTAATACTGCGGAACTGACTAAAGTAGT

PSC101 origin NNNNN; Patagonian FLP NNNNN; Chloramphenicol resistance cassette NNNNN

Target plasmid monomer:

TTTCCATAGGCTCCGCCCCCCTGACGAGCATCACAAAAATCGACGCTCAAGTCAGAGGTGGCGAAACCC
GACAGGACTATAAAGATACCAGGCGTTTCCCCCTGGAAGCTCCCTCGTGCGCTCTCCTGTTCCGACCCT
GCCGCTTACCGGATACCTGTCCGCCTTTCTCCCTTCGGGAAGCGTGGCGCTTTCTCATAGCTCACGCTG
TAGGTATCTCAGTTCGGTGTAGGTCGTTCGCTCCAAGCTGGGCTGTGTGCACGAACCCCCCGTTCAGCC
CGACCGCTGCGCCTTATCCGGTAACTATCGTCTTGAGTCCAACCCGGTAAGACACGACTTATCGCCACT
GGCAGCAGCCACTGGTAACAGGATTAGCAGAGCGAGGTATGTAGGCGGTGCTACAGAGTTCTTGAAGTG
GTGGCCTAACTACGGCTACACTAGAAGGACAGTATTTGGTATCTGCGCTCTGCTGAAGCCAGTTACCTT
CGGAAAAAGAGTTGGTAGCTCTTGATCCGGCAAACAAACCACCGCTGGTAGCGGTGGTTTTTTTGTTTG
CAAGCAGCAGATTACGCGCAGAAAAAAAGGATCTCAAGAAGATCCTTTGATCTTTTCTACGGGGTCTGA
CGCTCAGTGGGTCGACGTCTTACTGTCCCTAGTGCTTGGATTCTCACCAATAAAAAACGCCCGGCGGCA
ACCGAGCGTTCTGAACAAATCCAGATGGAGTTCTGAGGTCATTACTGGATCTATCAACAGGAGTCCAAG
CGAGCTCTCGAACCCCAGAGTCCCGCTCAGAAGAACTCGTCAAGAAGGCGATAGAAGGCGATGCGCTGC
GAATCGGGAGCGGCGATACCGTAAAGCACGAGGAAGCGGTCAGCCCATTCGCCGCCAAGCTCTTCAGCA
ATATCACGGGTAGCCAACGCTATGTCCTGATAGCGGTCCGCCACACCCAGCCGGCCACAGTCGATGAAT
CCAGAAAAGCGGCCATTTTCCACCATGATATTCGGCAAGCAGGCATCGCCATGGGTCACGACGAGATCC
TCGCCGTCGGGCATGCGCGCCTTGAGCCTGGCGAACAGTTCGGCTGGCGCGAGCCCCTGATGCTCTTCG
TCCAGATCATCCTGATCGACAAGACCGGCTTCCATCCGAGTACGTGCTCGCTCGATGCGATGTTTCGCT
TGGTGGTCGAATGGGCAGGTAGCCGGATCAAGCGTATGCAGCCGCCGCATTGCATCAGCCATGATGGAT
ACTTTCTCGGCAGGAGCAAGGTGAGATGACAGGAGATCCTGCCCCGGCACTTCGCCCAATAGCAGCCAG
TCCCTTCCCGCTTCAGTGACAACGTCGAGCACAGCTGCGCAAGGAACGCCCGTCGTGGCCAGCCACGAT
AGCCGCGCTGCCTCGTCCTGCAGTTCATTCAGGGCACCGGACAGGTCGGTCTTGACAAAAAGAACCGGG
CGCCCCTGCGCTGACAGCCGGAACACGGCGGCATCAGAGCAGCCGATTGTCTGTTGTGCCCAGTCATAG
CCGAATAGCCTCTCCACCCAAGCGGCCGGAGAACCTGCGTGCAATCCATCTTGTTCAATCATGCGAAAC
GATCCTCATCCTGTCTCTTGATCAGATCTTGATCCCCTGCGCCATCAGATCCTTGGCGGCAAGAAAGCC
ATCCAGTTTACTTTGCAGGGCTTCCCAACCTTACCAGAGGGCGCCTGACTGGAGTTCAGACGTGTGCTC
TTCCGATCTGTGGGTACCTTGTTGACTAGTGTGAGATCGGAAGAGCGTCGTGTAGGGAAAGAGTGTCCA
GCTGGCAATTCCGTTAATTAACACCTGACGTCTAAGAAACCATTATTATCATGACATTAACCTATAAAA
ATAGGCGTATCACGAGGCCCTTTCGTCTTCAAGAAGTTCCTATTCTCTAGAAAGTATAGGAACTTCTCA
CCACAGCTAACACCACGTCGTCCCTATCTGCTGCCCTAGGTCTATGAGTGGTTGCTGGATAAC**~Promo
ter~**ATATTCAGGGAGACCACAACGGTTTCCCTCTACAAATAATTTTGTTTAACTTTTCTAGATTTAAG
AAGGAGATATACAT**~Insert~**TAAATGTCCAGACCTGCAGGCATGCAAGCTCTAGAGGCATCAAATAA
AACGAAAGGCTCAGTCGAAAGACTGGGCCTTTCGTTTTATCTGTTGAAATGCACCAAAAACTCGTAAAA
GCTCTGATGTATCTATCTTTTTTACACCGTTTTCATCTGTGCATATGGACAGTTTTCCCT**~Restrict
ionAssemblySite~**GCTAGCCTCGGGCAGCGTTGGGTCCTGGCCACGGGTGCGCATGATCGTGCTCC
TGTCGTTGAGGACCCGGCTAGGCTGGCGGGGTTGCCTTACTGGTTAGCAGAATGAATCACCGATACGCG
AGCGAACGTGAAGCGACTGCTGCTGCAAAACGTCTGCGACCTGAGCAACAACATGAATGGTCTTCGGTT
TCCGTGTTTCGTAAAGTCTGGAAACGCGGAAGTCAGCGCCCTGCACCATTATGTTCCGGATCTGCATCG
CAGGATGCTGCTGGCTACCCTGTGGAACACCTACATCTGTATTAACGAAGCGCTGGCATTGACCCTGAG
TGATTTTTCTCTGGTCCCGCCGCATCCATACCGCCAGTTGTTTACCCTCACAACGTTCCAGTAACCGGG
CATGTTCATCATCAGTAACCCGTATCGTGAGCATCCTCTCTCGTTTCATCGGTATCATTACCCCCATGA

ACAGAAATCCCCCTTACACGGAGGCATCAGTGACCAAACAGGAAAAAACCGCCCTTAACATGGCCCGCT
TTATCAGAAGCCAGACATTAACGCTTCTGGAGAAACTCAACGAGCTGGACGCGGATGAACAGGCAGACA
TCTGTGAATCGCTTCACGACCACGCTGATGAGCTTTACCGCAGCTGCCTCGCGCGTTTCGGTGATGACG
GTGAAAACCTCTGACACATGCAGCTCCCGGAGACGGTCACAGCTTGTCTGTAAGCGGATGCCGGGAGCA
GACAAGCCCGTCAGGGCGCGTCAGCGGGTGTTGGCGGGTGTCGGGGCGCAGCCATGACCCAGTCACGTA
GCGATAGCGGAGTGTATACTGGCTTAACTATGCGGCATCAGAGCAGATTGTACTGAGAGTGCACCATAT
GCGGTGTGAAATACCGCACAGATGCGTAAGGAGAAAATACCGCATCAGGCGCTCTTCCGCTTCCTCGCT
CACTGACTCGCTGCGCTCGGTCGTTCGGCTGCGGCGAGCGGTATCAGCTCACTCAAAGGCGGTAATACG
GTTATCCACAGAATCAGGGGATAACGCAGGAAAGAACATGTGAGCAAAAGGCCAGCAAAAGGCCAGGAA
CCGTAAAAAGGCCGCGTTGCTGGCGTT

PBR322 origin NNNNN; FRT site NNNNN; Kanamycin resistance cassette NNNNN; Barcode region NNNNN

Promoters:
ProA
**TTTACG**GGCATGCATAAGGCTCG**TAGGCT**
ProC
**TTTACG**GGCATGCATAAGGCTCG**TATGAT**

Restriction assembly site for mScarlet-I plasmid:
**GGCGCGCC**GAGAGG**GGATCC**
Restriction assembly site for mWatermelon plasmid:
**GGATCC**GAGAGG**GGCGCGCC**

Inserts:
**mScarlet-I**
ATGAGTAAAGGAGAAGCTGTTATTAAAGAGTTCATGCGCTTCAAAGTTCACATGGAGGGTTCTATGAAC
GGTCACGAGTTCGAGATCGAAGGCGAAGGCGAGGGCCGTCCGTATGAAGGCACCCAGACCGCCAAACTG
AAAGTGACTAAAGGCGGCCCGCTGCCTTTTTCCTGGGACATCCTGAGCCCGCAATTTATGTACGGTTCT
AGGGCGTTCATCAAACACCCAGCGGATATCCCGGACTATTATAAGCAGTCTTTTCCGGAAGGTTTCAAG
TGGGAACGCGTAATGAATTTTGAAGATGGTGGTGCCGTGACCGTCACTCAGGACACCTCCCTGGAGGAT
GGCACCCTGATCTATAAAGTTAAACTGCGTGGTACTAATTTTCCACCTGATGGCCCGGTGATGCAGAAA
AAGACGATGGGTTGGGAGGCGTCTACCGAACGCTTGTATCCGGAAGATGGTGTGCTGAAAGGCGACATT
AAAATGGCCCTGCGCCTGAAAGATGGCGGCCGCTATCTGGCTGACTTCAAAACCACGTACAAAGCCAAG
AAACCTGTGCAGATGCCTGGCGCGTACAATGTGGACCGCAAACTGGACATCACCTCTCATAATGAAGAT
TATACGGTGGTAGAGCAATATGAGCGCTCCGAGGGTCGTCATTCTACCGGTGGCATGGATGAACTATAC
AAA
**mWatermelon**
ATGAGTAAAGGAGAAGCTCTGATTAAAGAGTACATGCGCTTCAAAGTTCACATGGAGGGTTCTATGGAC
GGTCACGAGTTCGAGATCGAAGGCGAAGGCGAGGGCCGTCCGTATGAAGGCACCCATACCGCCAAACTG
AAAGTGACTAAAGGCGGCCCGCTGCCTTTTTCCTGGGACATCCTGAGCCCGCAATTTGGCTACGGTTCT
AGGGCGTTCATCAAACACCCAGCGGATATCCCGGACTATTATAAGCAGTCTTTTCCGGAAGGTTTCAAG
TGGGAACGCGTAATGAATTTTGAAGATGGTGGTGCCGTGACCGTCACTCAGGACACCTCCCTGGAGGAT
GGCACCCTGATCCATAAAGTTAAACTGCGTGGTACTAATTTTCCACCTGATGGCCCGGTGATGCAGCGT

AAGACGATGGGTTGGGAGGCGTCTACCGAACGCTTGTATCCGGAAGATGGTGTGCTGAAAGGCGACATT
AAAATGGCCCTGCGCCTGAAAGATGGCGGCCGCTATCTGGCTGACTGCAAAACCACGTACAAAGCCAAG
AAACCTGTGCAGATGCCTGGCGCGTACAATGTGGACCGCAAACTGGACATCACCTCTCATAATGAAGAT
TATACGGTGGTAGAGCAATATGAGCGCTCCGAGGGTCGTCATTCTACCGGTGGCATGGATGAACTATAC
AAA

**mWatermelon-DfrA**

ATGAGTAAAGGAGAAGCTCTGATTAAAGAGTACATGCGCTTCAAAGTTCACATGGAGGGTTCTATGGAC
GGTCACGAGTTCGAGATCGAAGGCGAAGGCGAGGGCCGTCCGTATGAAGGCACCCATACCGCCAAACTG
AAAGTGACTAAAGGCGGCCCGCTGCCTTTTTCCTGGGACATCCTGAGCCCGCAATTTGGCTACGGTTCT
AGGGCGTTCATCAAACACCCAGCGGATATCCCGGACTATTATAAGCAGTCTTTTCCGGAAGGTTTCAAG
TGGGAACGCGTAATGAATTTTGAAGATGGTGGTGCCGTGACCGTCACTCAGGACACCTCCCTGGAGGAT
GGCACCCTGATCCATAAAGTTAAACTGCGTGGTACTAATTTTCCACCTGATGGCCCGGTGATGCAGCGT
AAGACGATGGGTTGGGAGGCGTCTACCGAACGCTTGTATCCGGAAGATGGTGTGCTGAAAGGCGACATT
AAAATGGCCCTGCGCCTGAAAGATGGCGGCCGCTATCTGGCTGACTGCAAAACCACGTACAAAGCCAAG
AAACCTGTGCAGATGCCTGGCGCGTACAATGTGGACCGCAAACTGGACATCACCTCTCATAATGAAGAT
TATACGGTGGTAGAGCAATATGAGCGCTCCGAGGGTCGTCATTCTACCGGTGGCATGGATGAACTATAC
AAATAAATGTCCAGACCTGCAGGCAGGGTGCGGCGGGCACACCGCCTCCCCTGAGCTGTCACCGGATGT
GCTTTCCGGTCTGATGAGTCCGTGAGGACGAAACAGCCTCTACAAATAATTTTGTTTAA**~RBS~**ATGAA
ACTATCACTAATGGTAGCTATATCGAAGAATGGAGTTATCGGGAATGGCCCTGATATTCCATGGAGTGC
CAAAGGTGAACAGCTCCTGTTTAAAGCTATTACCTATAACCAATGGCTGTTGGTTGGACGCAAGACTTT
TGAATCAATGGGAGCATTACCCAACCGAAAGTATGCGGTCGTAACACGTTCAAGTTTTACATCTGACAA
TGAGAACGTATTGATCTTTCCATCAATTAAAGATGCTTTAACCAACCTAAAGAAAATAACGGATCATGT
CATTGTTTCAGGTGGTGGGGAGATATACAAAAGCCTGATCGATCAAGTAGATACACTACATATATCTAC
AATAGACATCGAGCCGGAAGGTGATGTTTACTTTCCTGAAATCCCCAGCAATTTTAGGCCAGTTTTTAC
CCAAGACTTCGCCTCTAACATAAATTATAGTTACCAAATCTGGCAAAAGGGT

DfrA NNNNN

Ribosomal binding sites

Strong:

AAGGAAATAAGGAGCTGTAGGAT

Weak:

GTATAATCGGCTAGTTCATAGTCGTT

DAM methylation mutations at origin

Original sequence:

AGCGTCAGACCACGTAGAAAA**GATC**AAAG**GATC**TTCTTGA**GATC**CTTTTTTTCTGCGCGTAA

New sequence:

AGCGTCAGACCACGTAGAAAA**GATT**AAAG**GGTC**TTCTTGA**GATC**CTTTTTTTCTGCGCGTAA

Primers for barcode amplification

5’:

**AATGATACGGCGACCACCGAGATCTACACNNNNN**ACACTCTTTCCCTACACGAC

3':

**CAAGCAGAAGACGGCATACGAGATNNNNNGTGACTGGAGTTCAGACGTG**

Timestamp barcode **NNNNN**
